# Nucleolar reorganization on stress depends on physicochemical changes due to nascent rRNA synthesis

**DOI:** 10.64898/2026.01.03.697456

**Authors:** Sinjini Ghosh, Aprotim Mazumder

## Abstract

Successive maturation of ribosomal subunits occurs through multilayered phase-separated structures of the cell nucleolus. The spatio-functional relationship between transcription of rRNA and nucleolar substructures and how this adapts to cellular stress remain incompletely understood. In this study, we resolve the sub nucleolar structures using expansion microscopy to reveal ordered structures of fibrillar center (FC) and dense fibrillar component (DFC) domains as nested shells, which is reorganized upon cellular stress like DNA damage or RNA polymerase I (RNAPI) inhibition. Direct visualization of nascent (5’ ETS) and mature (28S) rRNA suggested that rRNA synthesis is the critical regulator of nucleolar size, and organization. Nucleolar reorganization upon stress emerges to be a direct function of nascent rRNA levels. Stress-induced transcription inhibition remodels the sub-nucleolar compartments from a viscoelastic state into solid-like condensates thereby perturbing the nucleolar pH gradient due to the missing rRNA scaffold. We show, that rather than signaling to mediate rDNA repair, nucleolar reorganization naturally arises primarily from reduced rRNA levels and the resultant biophysical restructuring of the nucleolus under cellular stress.

## Introduction

The nucleolus is the central site for ribosome production in eukaryotic cells, and nucleolar function is influenced by its structural organization. A key step in ribosome assembly in human cells begins with RNA polymerase I (RNAPI) synthesizing a 47S precursor rRNA transcripts from rDNA gene regions that is processed into three different rRNA (18S, 5.8S, 28S) (Pederson, 1998). These pre-rRNA molecules undergo multiple modification steps and then assemble with ribonucleoparticles (RNPs) to from the large and small ribosomal subunits (LSU, SSU) of the ribosome. rRNA synthesis is tightly regulated and is sensitive to conditions of stress and diseases. Research using diverse tools of electron and light microscopy, molecular biology and biochemistry, has led to a model where different sub-nucleolar domains correspond to different stages in ribosome biogenesis(Hernandez-Verdun, 2006a). rRNA synthesis occurs at the boundary between the fibrillar center (FC) and the dense fibrillar component (DFC) of the nucleolus, while later stages of ribosome maturation happen within the DFC and granular component (GC), respectively(Louvet *et al*, 2005). Key proteins in this process include the multi-HMG-box chromatin protein Upstream Binding Transcription Factor (UBF) residing in the FC, Fibrillarin (FBL) in the DFC and Nucleolin (NCL) in the GC. Recent studies have also revealed a fourth compartment termed as the nucleolar rim which houses proteins with higher intrinsic disorder (like Ki-67), but its function is still under investigation(Stenström *et al*, 2020). Under conditions of DNA damage or transcriptional stress, the three major compartments reorganize into a bipartite arrangement: in this conformation, the innermost FC and DFC compartments shift to the periphery of the nucleolus(Al-Baker *et al*, 2004). Another structural organization called the ‘nucleolar necklace’ can form under conditions of RNA polymerase II (RNAPII) elongation inhibition (treatment with 5, 6, dichloro-1-𝛽-D-ribofuranosylbenzimidazole (DRB)) wherein the FC-DFC modules remain intact while the granular compartment fragments into smaller masses(Louvet *et al*., 2005). Despite these insights, the detailed spatial arrangement of the nucleolus has only recently been investigated, with studies probing its ultrastructure to gain a better understanding of its tripartite organization(Maiser *et al*, 2020; Wei *et al*, 2024; Yao *et al*, 2019). However, the effects of cellular stress or disrupted transcription on nucleolar protein and RNA organization are still not well understood, limited both by the resolution of light microscopy, and lack of tools for detection of transcript species of interest and complex sample preparation in electron microscopy.

Of the many biological condensates within the cells like stress granules, Cajal bodies, paraspeckles and P-bodies and others, the multi-phase nucleolus is perhaps the most apparent and best studied(Benjamin R. Sabari1, 2020). The intrinsically disordered regions within the nucleolar proteins aid in its condensation into multilayered droplets while RNA binding domains of these proteins and rRNA itself can add to sub-compartment specificity(Carvalho *et al*, 2021; Lafontaine *et al*, 2021). The rRNA synthesized at the nucleolar core acts as a scaffold to segregate proteins into different compartments based on their function in the assembly line of making mature rRNA as parts of ribosomes(Riback *et al*, 2023; Wadsworth *et al*, 2024). rRNA enables multimodal interactions with proteins, while allowing immiscibility and stability among the sub-compartments due to its polyanionic nature(Berry *et al*, 2015; Lafontaine *et al*., 2021). The outflux of newly synthesized rRNA is facilitated by a combined action of K-block and D/E tracts enriched nucleolar proteins(Rai *et al*, 2023). The K-blocks create a scaffold of transient multivalent interactions that aid in nucleolar organization, while the D/E tracts serve as proton carriers that set up a pH gradient, both contributing to the directional export of processed rRNA(King *et al*, 2024). Additionally, the gradient of rRNA concentration and length provides enhanced stable differentiation among the sub-compartment owing to its surfactant-like properties(Yamamoto *et al*, 2023). But how the rRNA outward flux is spatially altered within the nucleolar sub-compartments in healthy versus stressed cells is still elusive, especially owing to high rRNA concentration within the sub-organelle, and diffraction limit of the optical systems. Recent super-resolution methods have changed this landscape(Correll *et al*, 2024; Maiser *et al*., 2020; Shan *et al*, 2023; Yao *et al*., 2019), and among these methods is expansion microscopy. This method is an emerging set of sample preparation techniques that involve embedding a sample in an expansible polymer and crosslinking its components to this polymer network(Asano *et al*, 2018; Cho *et al*, 2018; Gao *et al*, 2017). The polymer is then expanded in water, causing both the polymer and specimen to expand isotropically. This pulls the biomolecules apart enabling nanoscale imaging with diffraction limited optics thus bypassing the optical limitations of conventional light microscopes (Figure 1A). This tool has proven effective for analyzing a range of cellular structures across various biological specimens, including cultured cells, unicellular organisms, and tissues(Asano *et al*., 2018; Cahoon *et al*, 2017; Cho *et al*., 2018; Gao *et al*., 2017; Halpern *et al*, 2017; Wassie *et al*, 2019). However, its effectiveness for isotropic expansion and detailed study of a complex multicomponent compartment like the nucleolus is only beginning to get established. Recent studies have employed this method to the study of the chromatin or the nucleolus(Katelyn R. Alley, 2025; Pownall *et al*, 2023), but the biological structure-function relationship in the nucleolus and its stress-dependent reorganization with visualization of specific rRNA species remain unexplored.

**Figure 1:**
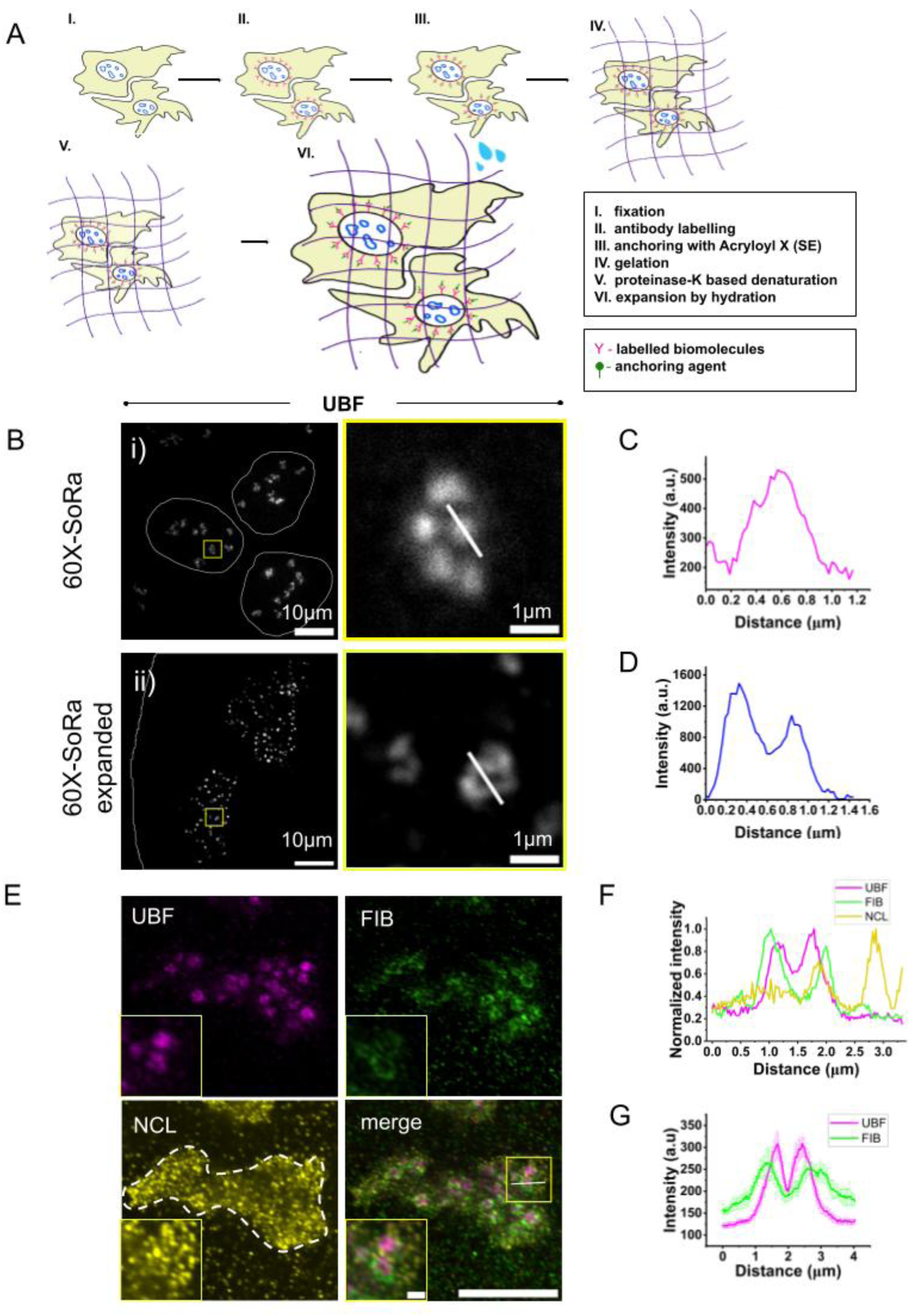
Expansion microscopy shows ultrastructure of the nucleolus comprising of core-shell sub-compartments: (A) Schematic for expansion microscopy. The cells are fixed after culturing on a coverslip (I), and then stained with antibody of choice (II). The cellular biomolecules are then anchored using Acryloyl X-SE (III), followed by gelation (IV), denaturation (V), and finally expansion of the gel by hydration. (B) Pre-expansion (top panel) and post expansion (bottom panel) image of U2OS cells stained with antibody against UBF1 taken in 60X SoRa, followed by inset from each of the images (bottom panel). Scale bar is 10 microns for expanded and 1 micron for unexpanded images. (C) Line profile for unexpanded (blue) and (D) expanded (magenta) UBF foci, marked by white lines in the top and bottom panel of (B) respectively. (E) Immunofluorescence assay performed on U2OS cells combined with expansion microscopy to mark all the three sub compartments of a single nucleolus: UBF (magenta; Fibrillar centres - FC), FIB (green; Dense fibrillar component - DFC), NCL (yellow; Granular component, GC), with an inset marked in yellow box highlighting 2-3 UBF-FIB rings. Scale bar is 10 microns for the nucleolus and 1 micron for the inset. (G) Single normalized intensity profile marked across all three sub-compartments shown by a white line in the merge image in (E). (H) The grand mean of intensity profiles from 20 FCs. For each FC, four radial lines were measured at 45° intervals and averaged to generate an individual profile. Data are presented as mean ± S.E.M (Standard error of mean) across the 10 individual FCs from a single experiment.

In this study, we utilized expansion microscopy in conjunction with super resolution by optical re-alignment (So-Ra) imaging to investigate the intricate details of the tripartite nucleolus under normal conditions and with cellular stress. Single-molecule Fluorescence in Situ Hybridization (smFISH) aids in the visualization of individual RNA molecules within a cell, achieved by using a set of multiple fluorescently-labeled oligomers designed against target RNA(Alley *et al*, 2025; Dhuppar & Mazumder, 2020; Femino *et al*, 1998; Pasnuri *et al*, 2023; Raj *et al*, 2008). We therefore, multiplexed immunofluorescence for nucleolar proteins with smFISH for rRNA in order to examine their relative spatial association. We assessed the correlation between the nascent rRNA synthesis with the spatial reorganization of the nucleolus under genotoxic or transcriptional stress. Such stress is well documented to cause nucleolar reorganization(Al-Baker *et al*., 2004; Boulon *et al*, 2010; Daniely *et al*, 2002a; Ljungman, 2000; Rubbi & Milner, 2003), but we show that nascent rRNA is a predictor of nucleolar reorganization. Stress-induced transcriptional downregulation in turn alters the physico-chemical properties of the nucleolus in terms rRNA and pH gradients, resulting in the realignment of the nucleolar sub-compartments under stress.

## Results

### Nucleolus comprises a core shell (FC-DFC) architecture immersed in a larger condensate of GC which reorganizes under stress as revealed by Expansion Microscopy

The nucleolus is the most distinct phase-separated organelle within the cell. But the detailed structure of the nucleolus along with different rRNA species have only recently started to be investigated in detail(Maiser *et al*., 2020; Yao *et al*., 2019). We used expansion microscopy to visualize the sub-domains of the nucleolus (Figure 1B-D) to gain more insight into nucleolar structure-function relationships. We first standardized the tool using the cell nucleus size as a readout using Protein-expansion microscopy (Pro-ExM)(Gao *et al*., 2017; Tillberg, 2016). We found that the ratio between the area of expanded nuclei to that of unexpanded nuclei resulted in an expansion factor of ∼4.18 (Supp. Figure 1A). This combined with So-Ra (a SIM-equivalent technology that improves lateral resolution by approximately a factor of 2 (Maiser *et al*., 2020)), which gives an additional 4X factor in 2D, we can effectively magnify the sample by ∼16X. The expansion factor not only serves to enlarge the sample, but also effectively improves resolution since the biomolecules that were in close proximity are now physically apart in space due to the expansion(Cho *et al*., 2018; Tillberg, 2016; Wassie *et al*., 2019) . The So-Ra provides an additional resolution of the sample by 1.4X (2X with deconvolution) which enables us to gain nanoscale insights on details that were previously underestimated due the diffraction limit of the microscope. Such expansion tools have been used for the study of chromatin and nucleolar structures recently, giving us additional confidence in our methods(Katelyn R. Alley, 2025; Pownall *et al*., 2023). We used multiplexed labelling of different nucleolar compartment proteins and observed them with So-Ra after expansion (Figure 1E). Expanded cells showed that both the DFC and FC components marked by Fibrillarin (FIB) and Upstream Binding Factor 1(UBF1) respectively, form ring-like structures where the FC ring is enclosed within the DFC ring (Figure 1F). The rings are oriented in X, Y and Z directions i.e. they form nested shells as observed in the 3D renders of FC-DFC compartments obtained using IMARIS (Supp. Figure 1B). We therefore quantified line profiles for the compartment markers over multiple such centers in different nucleoli to get average line profiles to conclude about the spatial relation of the concentric shells of UBF and FIB across cells (Figure1G). Considering the expansion factor during calculation, the FIB ringlets have an average diameter of 0.336 ±0.075 μm, in line with recent studies(Katelyn R. Alley, 2025; Maiser *et al*., 2020). We successfully further resolved the structure of UBF1 as the inner most ringlets (Figure1D) having an average diameter of 0.204 ±0.055 μm. While fibrillarin rings have been reported before(Lafontaine *et al*., 2021; Maiser *et al*., 2020; Yao *et al*., 2019), the small size of fibrillar centers make UBF rings hard to resolve. But using Structural Illumination Microscopy (SIM) imaging a recent study has reported UBF1 rings, giving us greater confidence about the structures we find(Maiser *et al*., 2020). We further corroborated the ring like organization of UBF1 using STED (stimulated emission depletion) microscopy, where UBF1 rings are further spatially resolved into clusters of 5-6 (single cross-sectional plane) beads in a single ring (Supp. Figure 1C). These multiple factories of nested FC-DFC exist in a bed of GC marked by Nucleolin (NCL) (Figure 1E). Moreover, we were also able to observe the nucleolar rim structures by staining for Ki-67 (Supp. Figure 2A) described in a very recent study(Stenström *et al*., 2020). We also look at other nucleolar proteins like RNAPI, nucleophosmin (NPM) to observe that they too reside in the FC and GC compartment respectively as expected (Supp. Figure 2A), but unlike UBF1, RNAPI does-not form a ring like structure as seen in recent literature as well(Katelyn R. Alley, 2025). Thus, the tool of expansion microscopy offers a unique handle to observe structural details of multiphase condensates of the tripartite nucleolus. A very recent study has indeed reported the use to expansion microscopy to study nucleolar structures(Alley *et al*., 2025), but beyond just reporting the structural details under normal homeostasis, we further investigated how it is reorganized under stress and the roles of rRNA in such reorganization.

**Figure 2:**
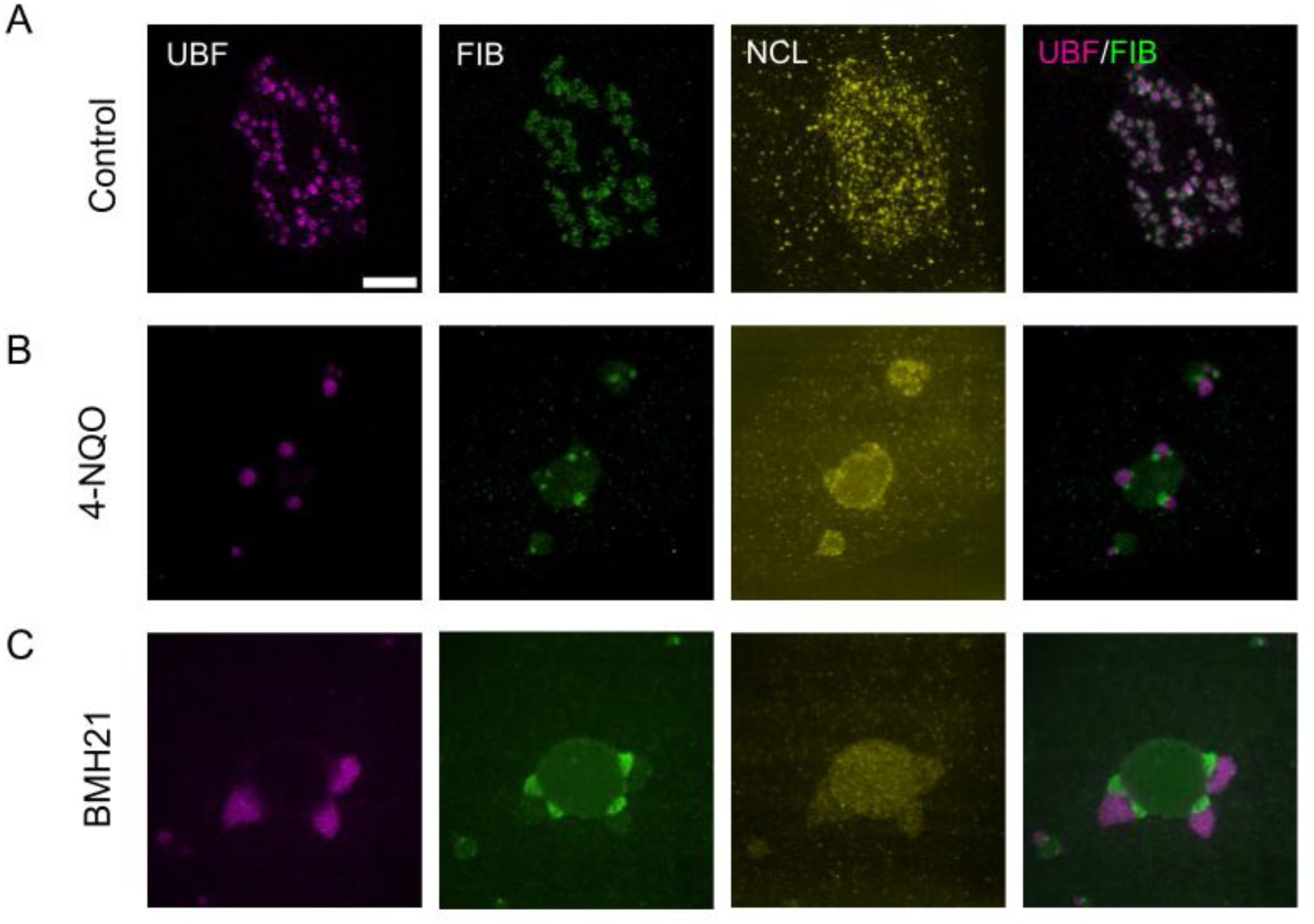
Expansion microscopy shows the reorganization of the nucleolus under genotoxic stress and transcription inhibition: (A) Representative expansion microscopy (ExM) image of a control nucleolus displaying tripartite organization. Nucleolar reorganization following treatment with (B) 1 μg/ml 4-NQO for 1 hr and (C) 2 μM BMH21 for 3 hrs. Samples used are U2OS cells immunostained for UBF (magenta), FIB (green), and NCL (yellow). Scale bar is 5 microns. Reorganization of UBF across the different stress conditions is shown later in Figure 5.

The spatial arrangement of the nested-shell emulsion structure of the nucleolus is a well-known biomarker for cell health, perturbation to which is seen in cancer and neurodegeneration(Carvalho *et al*., 2021; Dogra & Kriwacki, 2025; Penzo *et al*, 2019). Therefore, we investigated the impact of DNA damage and transcriptional inhibition on U2OS cells using 4-quinazoline-1-oxide (4-NQO, a UV-mimetic agent) and N-(2-(Dimethylamino) ethyl)-12-oxo-12H-benzo[g]pyrido [1-b] quinazoline-4-carboxamide (BMH21) (a RNAPI inhibitor) on nucleolar organization with expansion microscopy (Figure 2). Nucleolar cap formation due to DNA damage or transcriptional inhibition is known(Al-Baker *et al*., 2004; Boulon *et al*., 2010); but these studies are performed at diffraction-limited resolution and are further somewhat confounded by the fact that Actinomycin D used for RNAPI inhibition in some of these studies can itself trigger DNA damage responses(Mischo *et al*, 2005). The stress induced re-organization revealed that both UBF1 and FIB rings are transformed into coalesced single-phase condensates which re-arrange at the nucleolar periphery in an inside-out manner with UBF1 now on the outside. The UBF1 protrudes towards the nucleoplasm while FIB faces the inner nucleolar matrix of GC. NCL in this case shows lower intensity within the nucleoli (Figure 2A-C), suggesting its nucleoplasmic translocation for downstream repair of the damage as reported before(Daniely *et al*, 2002b; Kobayashi *et al*, 2012). The coalesced foci of UBF and FIB are parts of the ‘nucleolar caps’ known in earlier studies(van Sluis & McStay, 2017). But the nucleolus reorganizes differently based on how the transcriptional silencing occurs. If the noncoding RNAs (ncRNA) that help in the processing of the rRNA is blocked, the nucleolus segregates mainly affecting the DFC and GC compartment where the processing take place. Upon treating the cells with 5, 6, dichloro-1-𝛽-D-ribofuranosylbenzimidazole (DRB), a RNAPII mediated transcription elongation inhibitor, we observe that in contrast to RNAPI inhibition, the GC compartment (NPM, NCL) undergoes segregation forming broken beads of the ‘nucleolar necklace’ known from older studies (Supp. Fig 3A), while the FCs (UBF1) remains unaffected (Supp. Figure 3B). Although it is known that transcriptional inhibition and DNA damage leads to nucleolar reorganization(Boulon *et al*., 2010; Shav-Tal *et al*, 2005; Thomas Haaf, (1996); van Sluis & McStay, 2017), how the rRNA synthesis and maturation impact nucleolar phases and gate the outward rRNA flux is poorly understood. Some studies suggest that the damage to the DNA leads to nucleolus acting as a stress sensor to allow de-mixing of the condensate and its resident proteins to initiate repair, which leads to nucleolar segregation(Yang *et al*, 2018). Another idea is that damage to nuclear or nucleolar DNA eventually leads to reduced transcription that disrupts the RNA-protein interaction formed during active rDNA transcription. This leads to new interactions among proteins that eventually leads to protein rich heterotypic condensates that form the multiphase nucleolar caps(Korsholm *et al*, 2020). In order to understand the correlation among stress induced ribosomal biogenesis and nucleolar reorganization better, we next sought to visualize the structural details of rRNA within the cell.

**Figure 3:**
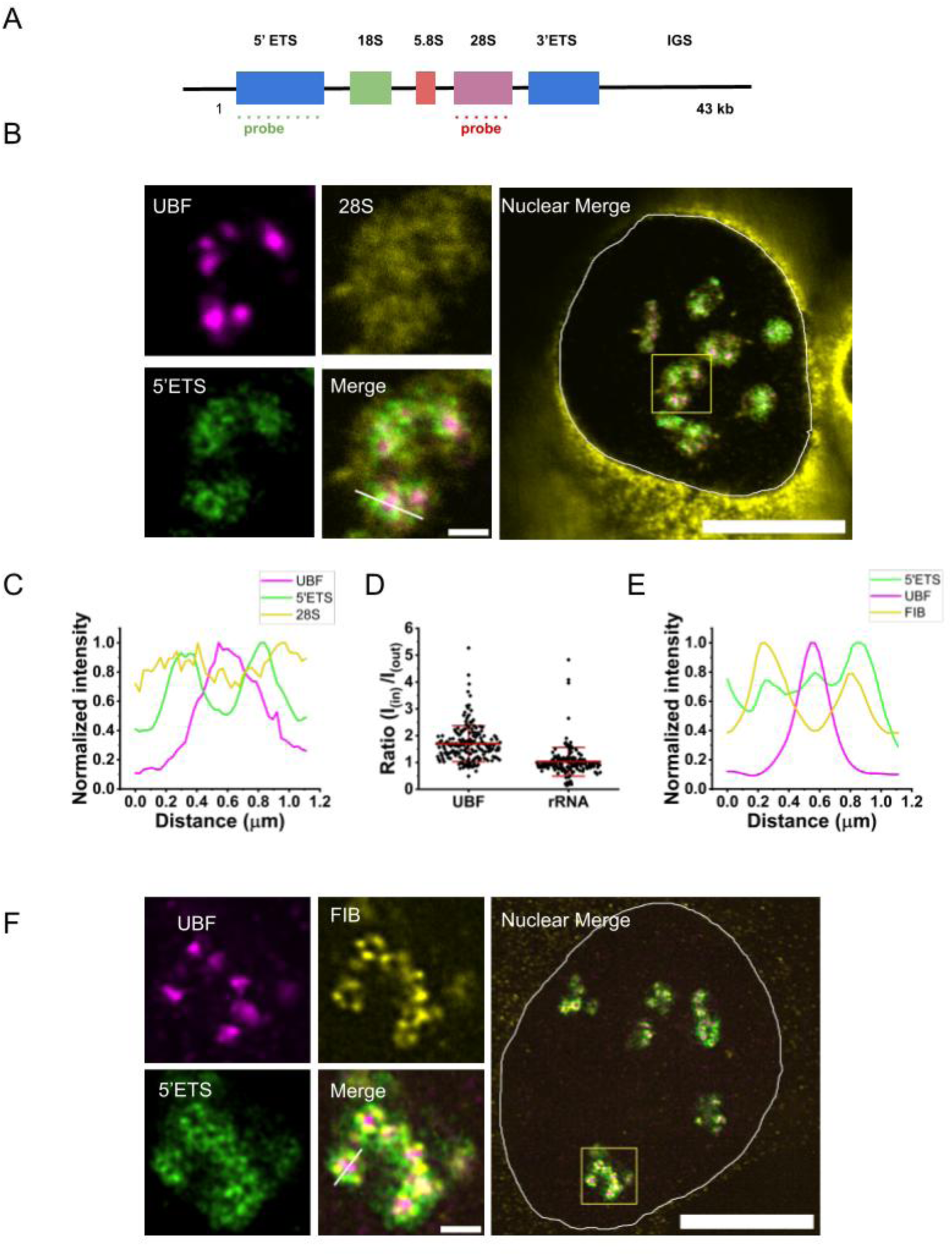
Nascent rRNA adds on to the nested multi-layered structure of the nucleolus: (A) Schematic for probes used in rRNA FISH. (B) Representative images from immunofluorescence assay on U2OS nucleoli stained with UBF (magenta), 28S mature rRNA(yellow) on the top panel, nascent 5’ETS rRNA (green) and the merge (bottom panel), with the nuclear merge on the right showing all the nucleoli in the given nucleus. Scale bar is 1 micron for the nucleoli and 10 microns for the nucleus. (C) Normalized intensity profile drawn across the merged inset (marked with white line), for UBF, 28S rRNA and 5’ETS rRNA. (D) Ratio of intensity of UBF and 5’ETS rRNA measured at the centre (I_in_) to the periphery (I_out_) of individual ring-shaped FC structures as seen in B, n = 21 nucleoli, over three independent replicates (E) Intensity profile of a line drawn across immunofluorescence images U2OS nucleoli shown in (F) marked with UBF (magenta). FIB (yellow) in top panel, 5’ETS (green) and the merge (bottom panel) with the nuclear merge on the right on the right showing all the nucleoli in the given nucleus. Scale bar is 1 micron for the nucleoli and 10 microns for the nucleus. Data represents mean± S.E.M. Statistical significance calculated using K.S. test (n.s is non-significant and *** represents p<0.0001)

### Nascent rRNA adds on to the nested multi-layered structure of the nucleolus which undergoes reorganization upon stress

rRNA molecules are co-transcribed as the 47S precursor rRNA(pre-rRNA) inside the nucleolus (approximately 13kb), flanked by the External Transcribed Spacers on both 5’ and 3’ end (5’ETS, 3’ETS), and interspersed by Internal Transcribed Spacer sequences (ITS1 and ITS2)(Aubert *et al*, 2018). The ETS get spliced by endonucleases during the earlier steps while the ITS get removed in later stages of processing and modifications(Aubert *et al*., 2018; Vanden Broeck & Klinge, 2024), to finally form the mature 28S, 18S, and 5.8S rRNA. The modified RNA then diffuses to GC where they assemble with the ribosomal subunits to form mature ribosomal subunits (Dogra & Kriwacki, 2025). The proteins aiding in this process of rRNA synthesis, processing and ribosomal assembly are organized into the nested multiphase nucleolus. Most of the earlier works have involved visualization of cellular RNA using RNA synthesis analogs like 5-ethynyl-uridine (5-EU) which mark the overall RNA pools in the cell(Jao & Salic, 2008). More recent studies have been investigating the structure of the nucleolus using rRNA sequence-specific probes by in-situ hybridization(Sofia A. Quinodoz *et al*, 2025; Wei *et al*., 2024; Yao *et al*., 2019). As rRNA represents the major pool of cellular RNA, EU does show enhanced signals in nucleoli, but it does not discriminate rRNA in different stages of processing. We find that though 5-EU showed some patterned structure inside the nucleoli (Supp. Figure 3C), but given this limitation, it is perhaps not the best tool to differentiate between nascent and mature rRNA transcripts. Therefore, to understand how the synthesized rRNA resides within the nucleolar structure, we designed probes for both nascent (5’ETS) and mature (28S) rRNA (Figure 3A) and used smFISH to identify their localization cell nucleoli. Given the high density of rRNA within nucleoli individual rRNA molecules are not distinguished like mRNA previous studies(Dhuppar & Mazumder, 2020; Pasnuri *et al*., 2023). But distinct spatial localization of 5’ETS and 28S were clear. Intensities of each species were measured at a single nucleolar level for quantification. We were unable to couple our smFISH protocol with expansion microscopy, and these experiments were performed with only So-Ra imaging, which stills offers improved resolutions by a factor of 1.4X over diffraction limited light microscopy. We observe that the mature rRNA marked by probes against 28S rRNA, are spread across the GC and the cytoplasm (where they are expected as a part of ribosomes), and excluded from the FCs. The FC-DFC boundary contains nascent rRNA which have not undergone splicing and modifications that are salient features of the mature rRNAs (Figure 3B). In contrast to the 28S mature rRNA signals, nascent rRNA marked by probes against the 5’ETS sequence, on the other hand, formed doughnut-like structures surrounding the FCs (marked by UBF1) (Figure 3B) as marked by the line profile (Figure 3C) drawn across the merged image in Figure 3B. Beyond just visual phenotypes, we quantified these ring-like structures by computing the ratio between the intensities at the center of the UBF1 foci to the outside. For a uniform distribution this enrichment metric should be 1. This enrichment metric is greater than 1 for 5’ETS suggesting it to be a hollow ring, while it is less than 1 for UBF1 itself as it marks the FC (Figure 3D). UBF1 rings are not apparent in this imaging mode without expansion and the protein just serves as a proxy for the FC. From our previous expansion microscopy-based experiments, we already know that UBF1 is in itself a 3D-shell like structure. Therefore, 5’ETS region is synthesized in an elongating bead-like manner on the ringlets of UBF1 forming a nested loop. Co-stained FC-DFC modules (using UBF1 and FIB) along with 5’ETS to understand the spatial localization of the nascent pre-rRNA showed that the pre-rRNA partially overlapped with FIB seen in the images as well as the line profile (Figure 3E, F). The partial interspersion of 5’ETS with FIB signals (Figure 3F) suggested that the rRNA transcription is initiated at the interface of FC-DFC with co-transcriptional processing, triggering phase separation and rRNA sorting into the subsequent compartments.

In-vitro studies have shown how RNA acts a scaffold for seeding liquid-liquid phase-separated (LLPS) condensates(Rai *et al*., 2023; Roden & Gladfelter, 2021), and it is known that reducing RNA levels can cause protein orders in condensates to be reversed in in-vitro experiments (Rai *et al*., 2023). It can also hasten the nucleation of the condensates by reducing the activation energy of the coalescence kinetics(Benjamin R. Sabari1, 2020; Lafontaine *et al*., 2021; Wadsworth *et al*., 2024). It is known that the rRNA leads to the nested tripartite structure of nucleoli(Farley *et al*, 2015; van Sluis & McStay, 2017) and direct or indirect inhibition of transcription leads to the nucleolar reorganization leading to its segregation or reoriented bipartite structure. So, we directly visualized the rRNA upon global DNA damage and transcriptional stress. We find that inhibition of RNAPI mediated transcription by BMH21 leads to complete abrogation of nascent rRNA visualized by 5’ETS probes (Figure 4A, B). Mature and processed rRNA marked by 28S rRNA probes, though significant, show modest changes compared to 5’ETS rRNA (Figure 4B, C). This is reasonable, because inhibition of transcription does not immediately degrade mature rRNA already synthesized. DNA damage by 4-NQO has a similar but less pronounced impact on the nascent rRNA. Correlations between nascent and mature rRNA levels were strong in control cells, and become markedly weak with cellular stress (Figure 4A). Cells administered with 4-NQO (1µg/ml, 1hour) revealed that even though nascent rRNA is not completely abrogated, it still leads to a bipartite rearrangement of FC-DFC as seen under the effects of transcriptional stress. However, the protrusion of UBF1 from the core of the nucleolus under these conditions was not as complete as with BMH21 (Figure 4A), providing an early hint that nascent rRNA levels may be a critical regulator of nucleolar reorganization. Nascent rRNA though present in this case, reorganizes to form larger rings (compared to control) with coalesced UBF foci at its center as compared to active FC ringlets seen in control cells with super resolution microscopy (Figure 2A, S1C). With higher downregulation of rRNA synthesis by BMH21, the residual pre-rRNA migrates to the nucleolar periphery forming a ring around the nucleolar cap. We also observe that the nucleolar shape becomes more circular concomitant with progressive decline in nascent rRNA levels (Figure 4D), and reduction in number of UBF foci (Figure 4E). Indeed, the shape of the nucleolus has been shown to be a marker of nucleolar activity(Farley-Barnes *et al*, 2018; Hernandez-Verdun, 2006b; Lafita-Navarro & Conacci-Sorrell, 2023). Additionally, nucleolar area gradually decreases with damage and complete abrogation of rRNA synthesis (Figure 4F), but the overall nucleolar numbers remain the same (Figure4G). Despite a reduction in size, the stability of nucleolar counts (Figure 4G) indicates a functional down shift rather than a structural disintegration. While the structural integrity of the individual Nucleolar Organizing Regions (NORs)-associated nucleolar assembly remains intact, their overall capacity of mature ribosomal production post rRNA synthesis is reduced under stress. Earlier reports suggested that the protein reorganization in both silencing of RNAPI-mediated transcription and DNA damage are similar(Franek *et al*, 2016), but we show that the spatial localization and levels of nascent rRNA can vary in case of DNA damage and transcriptional stress, perhaps related to the level of rRNA abrogation. To investigate this point further, we aimed to understand the relation between the spatial reorganization of the compartment-specific proteins and nascent rRNA levels with varying time after DNA damage.

**Figure 4:**
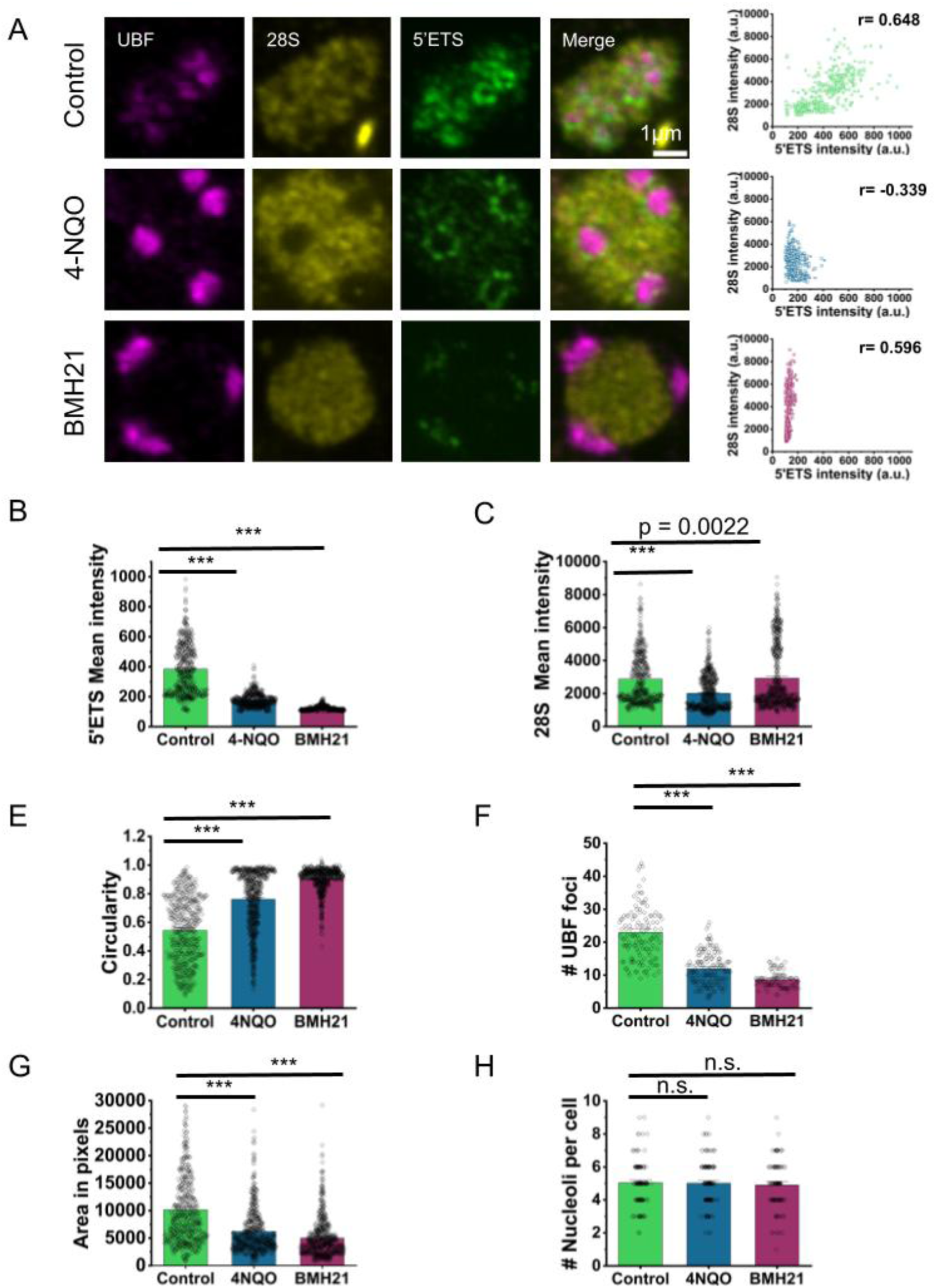
Nucleolus shows stress induced relocalization of its resident protein UBF and nascent rRNA: (A) Representative immunofluorescence images of U2OS cell nucleoli marked with UBF (magenta), 28S rRNA (yellow), 5’ETS (green), and its merge, and compared with the same of cells treated with 1ug/ml 4NQO for 1hr (middle panel) and 2uM of BMH21 for 3hrs (lower panel) with a correlation study between intensity levels of 28S rRNA and 5’ETS rRNA. (B-G) various nucleolar parameters calculated from the experiment in (A). (B) Mean intensity of 5’ETS in individual nucleoli, and (C) Mean intensity of 28S rRNA, n ≥ 459 nucleoli, (D) Circularity of nucleoli, n ≥ 568 nucleoli (E) Number of UBF foci, n ≥ 90 nucleoli (F) Nucleolar area, n ≥ 449 nucleoli and (G) Number of nucleoli per cell (n>100 nucleoli, from two independent replicates). Data from (B-F) represents mean± S.E.M. Statistical significance calculated using K.S. test (n.s is non-significant and *** represents p<0.0001). Scale bar is 1 micron. All data except (G) are from three independent experiments

### Cellular rRNA levels and organization scales with the stress induced spatial rearrangement of nucleolar compartments

The cell nucleolus is emerging as a stress sensor within the cells(Yang *et al*., 2018). It has gained popularity as a potential biomarker for pathological conditions of cancer and neurodegeneration in the past decade. But most of the studies have looked at rRNA using either bulk methods or 5-EU at long term damage intervals(Korsholm *et al*., 2020; Szaflarski *et al*, 2022). These data suggested rRNA synthesis is significantly downregulated along with nucleolar cap formation that leads to migration of the damaged loci to the nucleolar caps allowing access to repair factors earlier known to be excluded from the nucleolar interior(Franek *et al*., 2016; van Sluis & McStay, 2017). But how the rRNA localization is actually spatially altered along with reduced synthesis upon DNA damage is not well understood. Therefore, to see if persistent DNA damage can cause further abrogation of rRNA levels, we looked at an additional timepoint of 4-NQO damage (1µg/ml, for 3hours), comparing it to cells administered with same dose but lower time of DNA damage (1 hour) and transcriptional stress by BMH21 (as discussed in previous section). We found that the increased exposure to 4-NQO led to further migration of the FC-DFC modules towards the nucleolar periphery (Figure5A). The nascent rRNA is decorated at the nucleolar periphery with fewer rings of nascent rRNA enveloping the FC-DFC cap formed at the periphery, suggesting that increased damage time leads to further downregulation of nascent rRNA and more fusion of the protein condensates forming coalesced foci at the nucleolar perimeter. To quantify the nucleolar reorganization, we designed a metric where stress induced reorganization of UBF is quantified and correlated to the nascent rRNA levels. The nucleolar perimeter is marked and the nascent rRNA intensity within the nucleoli is computed. The FC centers marked by UBF1 in a nucleolus are segmented, and the centroid is calculated for each. The shortest distance between the individual UBF1 foci and the nucleolar periphery is calculated. Distance of each such UBF focus is calculated from the nucleolar perimeter, with positive values indicating foci within the nucleolar perimeter and negative values indicating foci whose centroid lay outside the nucleolar periphery, in effect indicating extrusion of the foci. The average distance of the UBF1 foci from the nucleolar periphery is then correlated to the changes in the intensity of the nascent rRNA levels, in a given nucleolus. The nascent rRNA show graded reduction in across control, 4NQO-1h, 4-NQO-3h, and BMH21 respectively, and it positively scales with the movement of FC compartments progressively towards the periphery with a Pearson’s correlation coefficient of 0.569 (Figure 4B). UBF1 and FIB coalesce into larger condensates upon decreasing rRNA synthesis. This is possibly due to greater homo and heterotypic protein-protein interaction in the absence of synthesizing rRNA as compared to RNA-protein interactions in actively transcribing population of cells(Lafontaine *et al*., 2021; Lian Zhua, 2019). Therefore, we next probed into the dynamic biophysical properties of the nucleolar sub-compartments that regulate the nucleolar assembly, to better understand how stress alters the spatial and chemical properties of the sub-nucleolar condensates, owing to their reorganization.

### The physicochemical properties of the condensates alter in response to DNA damage and repressed transcription, leading to the bipartite reorganization of the nucleolus

Condensates formed by RNA or protein or both, get seeded by upon increasing concentration and affinity towards each other(Berry *et al*., 2015; Brangwynne *et al*, 2011; Rai *et al*., 2023). These are altered in various stress conditions and even though the various condensates still remain, change in their composition can lead to perturbed dynamic properties of these biomolecular condensates. Therefore, we investigated the temporal dynamics of the FC-marker protein UBF1, which regulates active transcription within the nucleoli using tools of Fluorescent Recovery After Photobleaching (FRAP). Nucleoli in U2OS cells transiently expressing UBF-GFP(Dundr *et al*, 2002) were photobleached under normal conditions and upon both DNA damage (by 4-NQO) and transcriptional inhibition (by BMH21), using a 488 nm laser, and the recovery dynamics of the fluorescent fusion protein was captured for 3 minutes (Figure 5A). Previous studies have shown that treatment with Actinomycin-D can cause reduction in the FRAP recovery of GFP-UBF as compared to other GC proteins like NCL, NPM1 and ribosomal protein S5, suggesting change in the FC protein dynamics which reorganizes into caps upon RNAPI mediated transcription inhibition by Actinomycin-D(Chen & Huang, 2001). In our study, we observed a very modest effect on the GFP-UBF recovery upon DNA damage over 1 hour and 3 hours (half time :t1/2 changed from 13.59s ±0.336s in controls to 14.73s ±0.481s (1 h) or 16.46s ±0.566s (3h)) post 4-NQO incubation, while transcriptional inhibition (by BMH21) showed significant loss in recovery at the photobleached site (t1/2 of 43.77s) (Figure 5B, C). The region of photobleaching was mainly restricted to one entire nucleolus per cell wherein any recovery expected is mostly from the nucleoplasm. We also tried partial nucleolar FRAP but the result did not differ significantly indicating that irrespective of the area of bleaching, the translational dynamics of UBF1 is lowered upon RNAPI transcriptional inhibition. It is interesting to note that while both the DNA damage conditions and the RNAPI inhibition cause reorganization as evidenced by the flipping of the nucleolar compartments (Figure 3G, 4A), there is still residual rRNA transcription under both the 4NQO damage conditions used (Figure 4A); but once the reorganization is complete with total transcriptional inhibition by BMH21, the dynamics of UBF also changes in a pronounced manner (Figure 5A-C). In-vitro studies have shown that loss in recovery of biomolecular droplets suggest a shift in the phases of these condensates(Lafontaine *et al*., 2021; Lian Zhua, 2019; Sheu-Gruttadauria *et al*, 2024). Transcriptional inhibition is known to cause changes in the visco-elastic properties of nucleolus(Riback *et al*., 2023). Therefore, we infer that the loss in recovery of GFP-UBF indicates transition of these condensates to a low dynamics solid-like state compared to their original viscoelastic gel-like state when nucleolar transcription was active. Reduction in rRNA can cause a change in the molecular balance in the structures that form the signatures sub-compartments(Caragine *et al*, 2019; Farley *et al*., 2015; Lafontaine *et al*., 2021; Lawrimore *et al*, 2021). This is because rRNA can scaffold multivalent interaction among the Glycine-Arginine rich (GAR) resident proteins which, when altered, shifts to GAR-GAR interacting condensates making it more viscous or dense in nature(Feric *et al*, 2016; Riback *et al*., 2023).

**Figure 5:**
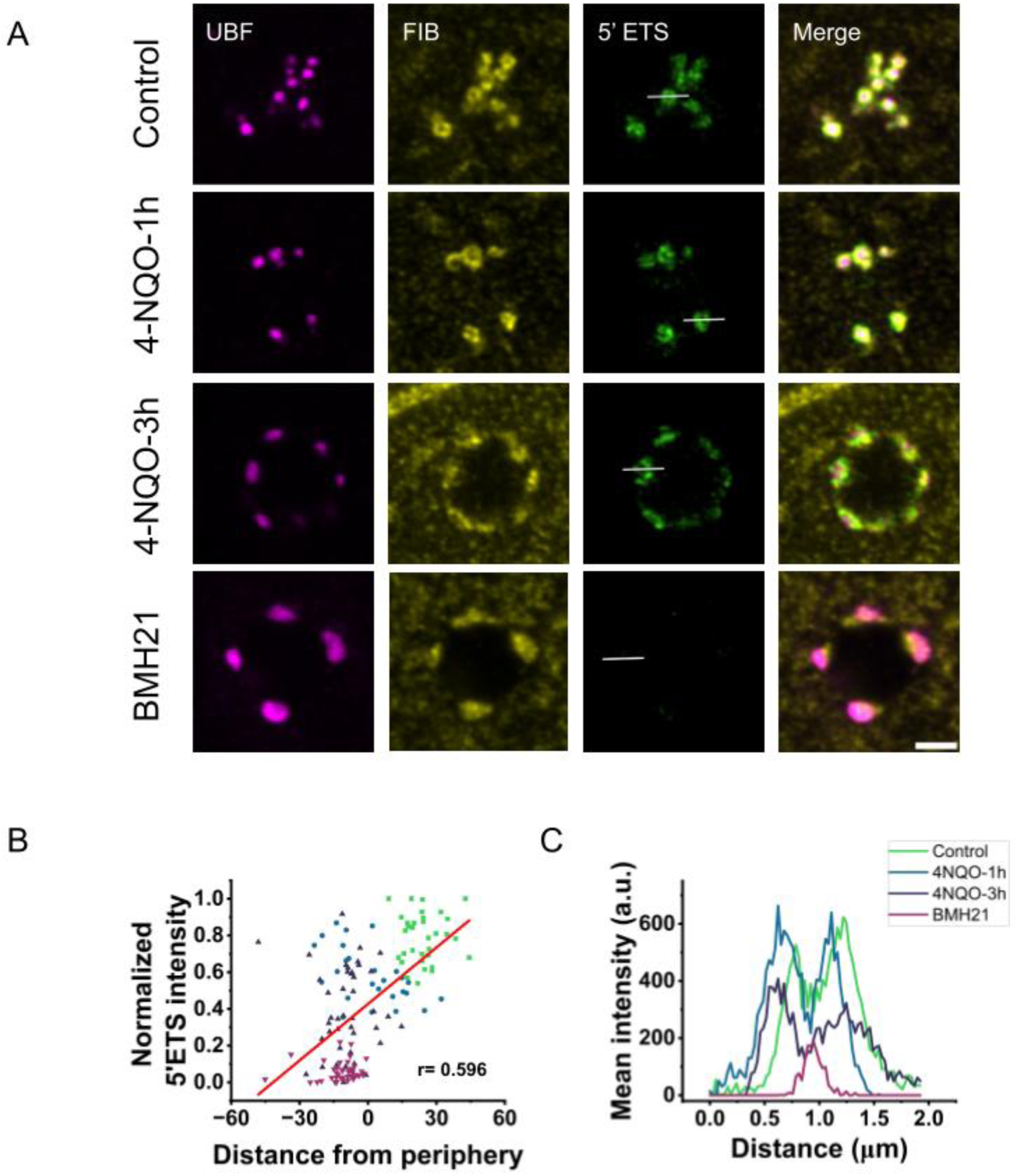
Genotoxic stress leads to similar nucleolar organization as seen in transcriptional inhibition by BMH21: (A) Representative images from immunofluorescence assay of U2OS cell nucleoli marked with UBF(magenta), FIB (yellow), 5’ETS (green), and its merge, and compared to that of cells treated with 1ug/ml 4NQO for 1hr, 3hr and 2uM of BMH21 for 3hrs. Scale bar is 1 microns (B) Scatterplot of distance of UBF foci from the nucleolar periphery, against the rRNA intensity of the respective nucleolus. Pearson’s correlation coefficient (r) calculated. N ≥ 26 nucleoli, across three independent experiments. (C) Spatial arrangement of rRNA across the images shown in (A) with a line profile. Scale bar is 1 micron.

The nucleolar proteins along with their GAR-rich domains also often contain Intrinsically Disordered Regions (IDRs) enriched in charged and polar amino-acids that promote multivalent, electrostatic, dipole-dipole and hydrogen bonding interactions with the newly synthesized rRNA(Dogra & Kriwacki, 2025). This facilitates an intrinsic pH gradient and help maintain different chemical microenvironments simultaneously within the cell. The nucleolus thus inherently shows a lower pH than the rest of the nucleoplasm(Aryan *et al*, 2023; King *et al*., 2024; Martin *et al*, 2015). To further understand how the nucleolar environments can be affected with morphological remodeling of the tripartite condensate, we tested the effect of DNA damage and transcriptional downregulation on the pH of the cell nucleoli. To do this, we used the cell permeant ratio-metric pH sensor Seminaphtharhodafluor 5-(and-6)-carboxylic acid, acetoxymethyl ester, acetate (SNARF-4F, AM ester), as done in previous studies(Aryan *et al*., 2023; King *et al*., 2024). This sensor when excited with 561nm emits in both 580 and 640nm range. The extent of protonation-deprotonation of the dye varies with cellular pH which is reflected in the emissions collected at the two emission wavelengths(Marcotte & Brouwer, 2005). The ratio of the emissions at the two wavelengths is a proxy for the pH recorded inside the cells. Higher 640/580 ratios indicate higher pH and thus deacidification of the cell or organelle of interest. We observed that the 640/580 ratio increases significantly upon DNA damage and more modestly but significantly with RNAPI inhibition (Figure 5D, E). Beyond just reducing rRNA levels, DNA damage can cause translocation of nucleolar proteins like NCL and NPM 1 (Daniely *et al*., 2002b; Yang *et al*, 2016), and also PARP1, which synthesizes negatively charged poly-ADP-ribose (PAR) chains essential for rRNA processing (Aryan *et al*., 2023; Boamah *et al*, 2012) potentially contributing to nucleolar acidification. NCL and NPM 1 are themselves rich negatively charged DE tracts (King *et al*., 2024; Mitrea *et al*, 2016; Scott & Oeffinger, 2016). Together, the data from immunofluorescence and smFISH (Figure 4A) and FRAP studies suggest that decline in rRNA synthesis can alter the structural organization and material properties of the nucleolus. Further in the context of DNA damage, shuttling of nucleolar proteins, potential loss of nucleolar PAR chains, in addition to reduced rRNA levels, may alters the molecular profile of the nucleolus potentially leading to a more pronounced nucleolar deacidification. The GFP-UBF FRAP (Figure 6A) and pH studies (Figure 6D, E), show that while there are common features, different stresses can alter the biophysical properties of sub-nucleolar condensates in more nuanced ways than just broad reorganization.

**Figure 6:**
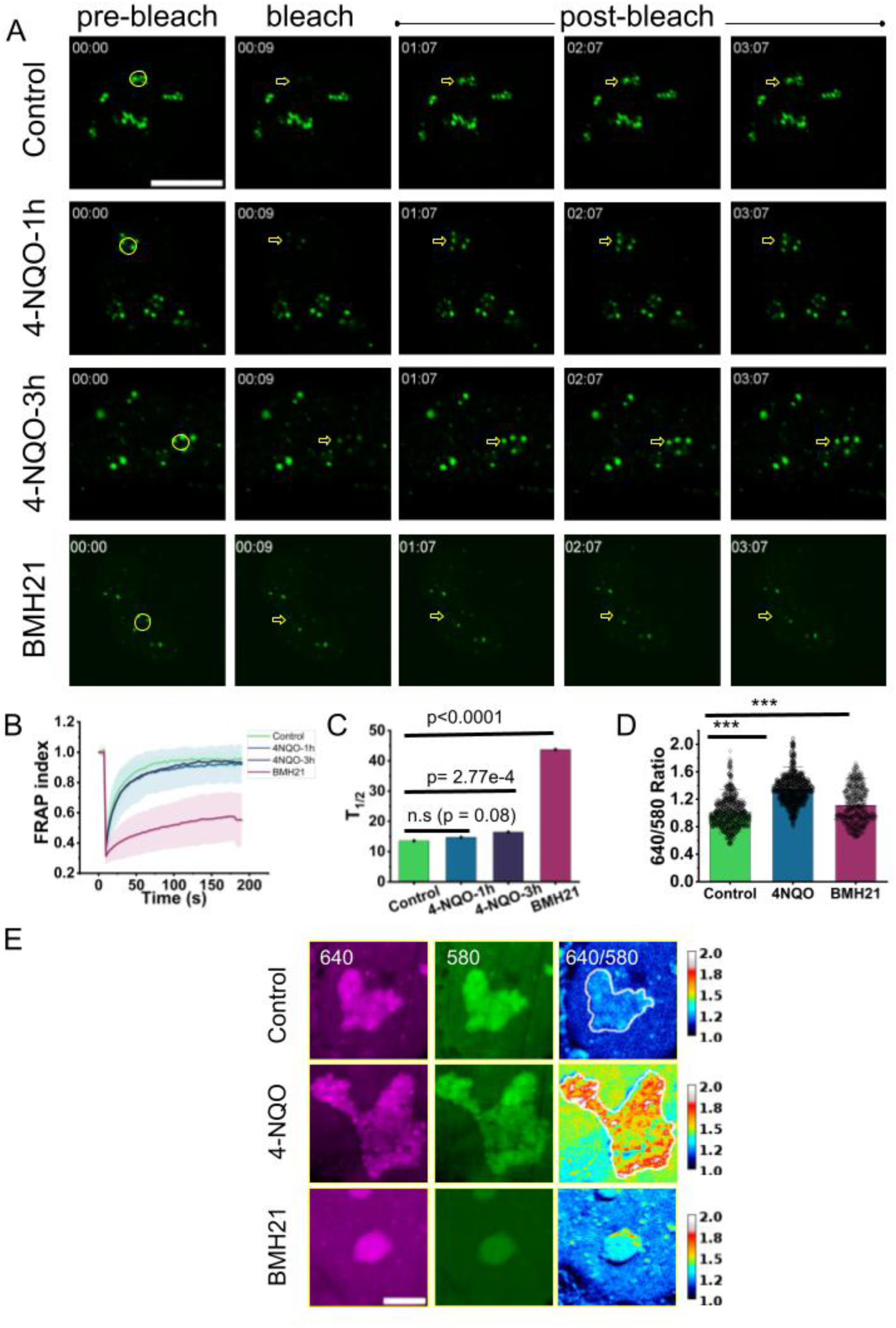
The biophysical properties of the condensates alter in response to repressed transcription, leading to the reorganized bipartite arrangement of the nucleolus: (A) Representative images of time-lapse of U2OS cells transiently transfected with GFP-UBF with a single nucleolus photobleached for every cell using 488 laser, in control, and compared with cells treated with 1ug/ml 4-NQO for 1hr and 3hrs, and 2uM BMH21 for 3hrs respectively, and (B) FRAP curve for the same experiment. Scale bar is 10 microns. Data represents mean± S.E.M. n>20 cells, from 3 independent experiments. (C) T-half (T_1/2_) calculated from (B), (D) 640/580 ratios calculated using the experiment from (E) which shows U2OS cells incubated with 7.5 um SNARF-4F for 30 min. The images show a single nucleolus from each treatment of 1ug/ml 4-NQO for 1 hr and 2uM BMH21 for 3 hrs respectively compared to the control cells. Scale bar is 5 microns. The samples were excited with 561 nm and emission was collected at 640 and 580 nm. Data represents mean± S.E.M, n>100, from 3 independent experiments. Statistical significance calculated using K.S. test (n.s is non-significant and *** represents p<0.0001)

**Figure 7:**
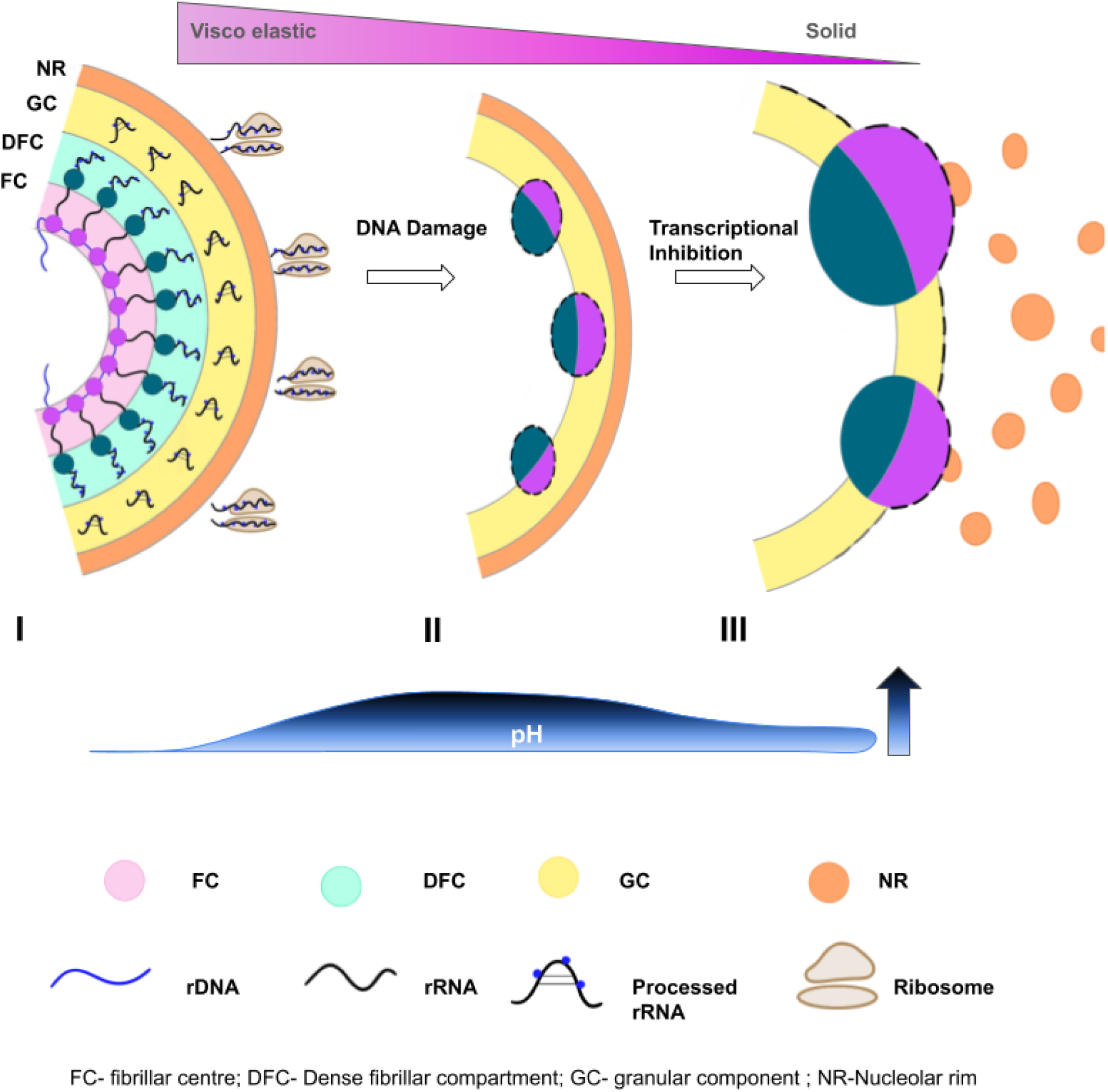
Schematic model for the damage induced nucleolar reorganization.: (I) In a healthy homeostatic state, the nucleolus is characterized by a liquid-like tripartite architecture comprising of multiple subdomains which aid in the transcription of rRNA for formation of mature ribosomes. The FCs (marked in light pink, with magenta marking UBF) and DFCs (light green, with FIB labelled in dark green) within the nucleolus forms ring-like structure embedded within a pool of granular component. A fourth compartment called the nucleolar rim form a ring around the nucleolar periphery. The rDNA (blue) is actively transcribed into nascent pre-rRNA (black) at the FC-DFC interface (which in itself looks like a nested ring within) and co-transcriptionally modified, forming mature ribosomes for directional export out into the nucleoplasm after assembly. This liquid-like state exhibits high molecular mobility and a balanced internal pH, maintained by rRNA molecules and charged disordered nucleolar proteins acting as essential proton carriers. Upon global DNA damage, such as exposure to 4-NQO (II), the rRNA production is attenuated, initiating a nucleolar reorganization that leads to formation of FC and DFC foci disrupting their ring like organization. The GC pool of proteins like NCL, which aid in repair processes shuttle outside the nucleoplasm. This leads to change in the pool of proteins within the nucleoli with missing scaffold of rRNA shifting the interaction paradigm within the proteins and rRNA which now no longer are available to serve as proton carriers, leading to increase in nucleolar pH. Further abrogation of rRNA mostly due to silencing of RNAPI mediated transcription leads to flipping out of the tripartite organization of the FC-DFC-GC into a bipartite structure. In this final stage of pronounced reduction in nascent rRNA, the nucleolus undergoes a phase transition to a solid-like state with diminished protein mobility. This is marked by the structural inversion, where the FCs completely moves to the nucleolar periphery facing outwards, whereas the FIB is at the periphery facing the nucleolar interior. The residual rRNA is also decorated at the nucleolar periphery. Ultimately, the model illustrates that a graded reduction in nascent rRNA levels dictates the transition of nucleolus from a dynamic biosynthetic hub to a rigid, segregated body with fundamentally altered physical and biochemical properties.

## Discussion

The nucleolus is perhaps the best-known phase-separated sub-organelle of the cell. Despite that, its dynamic organization inside the cell especially under conditions of environmental stress is underappreciated still, and recent studies have opened new avenues for studying this organelle with improved spatial resolution(Maiser *et al*., 2020; Yao *et al*., 2019). In this study we investigated nucleolar organization at a higher spatial resolution to understand its structure-function correlation better. We first used the tools of expansion microscopy to study the nucleolar sub-compartments (Figure 1), with specific resident proteins for the sub-compartments as a proxy to label them. We found that the innermost compartment protein, UBF1, itself shows a shell-like appearance with a pool of multiple such rRNA-making factories occupying the entire nucleolus. Shells of the DFC surrounded these and are embedded in the GC (Figure 1E). Nucleolar reorganization with DNA damage or transcriptional stress (Figure 2) could be followed beyond just simple cap formation. Shape and size parameters of nucleoli could be related to rRNA production activity. We further wanted to understand how nucleolar RNA is spatially arranged within the cell nucleolus. Probing the cells with both mature and nascent pre-rRNA (Figure 3) we find that rRNA is synthesized in the FC-DFC boundary and then gradually progresses to the GC potentially undergoing modifications and cleavage in line with previous studies (Yao *et al*., 2019). The mature rRNA then gets assembled at GC as seen by the homogenous staining of 28S rRNA probe across the entire GC with a small zone of low intensity which marks the core of the FC marked with UBF (Figure 3B). This corroborates that mature rRNA is mainly excluded from the FC. The nascent pre-rRNA on the other hand, forms an intermediate shell that partially overlaps with the DFC region marked by FIB (Figure 3F). The cells upon stress undergoes reorganization of the nucleolus that leads to a switch from the tripartite to the bipartite arrangement wherein the inner compartment flips outside the nucleolar periphery and the DFC forms a clustered condensate inwards. We observe this upon treatment with DNA-damaging 4-NQO and RNAPI transcription inhibiting BMH21 (Figure 2, 4A). Although the 28S rRNA levels showed divergent trends upon the aforementioned dosage of DNA damage, and RNAPI inhibition, its correlation with the nascent rRNA levels reduced dramatically with decreasing synthesis of pre-rRNA (Figure 4A). The nucleolar proteins show bipartite arrangement as seen by earlier studies(Al-Baker *et al*., 2004; Boulon *et al*., 2010). But what was interesting to note is that upon DNA damage with 4NQO, rRNA levels do not get abrogated entirely, and nascent rRNA levels are a predictor of nucleolar reorganization (Figure 4, 5). Other DNA damaging agents may cause different nucleolar reorganization phenotypes, correlated with rRNA level reductions. This remains to be explored. In our experiments, the reduction in rRNA synthesis leads to coalescence of the remaining FC-DFC modules, the size of the FC-DFC units increases, while the nucleolar number remains stable. This is most likely because, although the NOR-associated ‘factory sites’ prevail, their functional capacity to produce mature ribosomes is diminished due to stress. The prevailing nascent rRNA transcripts then decorate these coalesced foci initiating the cap formation, which is seen as rings of 5’ETS rRNA upon treatment with 4-NQO (Figure 4A, 5A). We designed a metric to study the nucleolar reorganization at the level of individual nucleoli across cells, based on the distance of the UBF1 foci from the nucleolar periphery. We simultaneously measure the nucleolar nascent rRNA levels in the same nucleoli. Nucleolar reorganization emerges to be strongly correlated to rRNA levels (Figure 5B). Persisting damage for longer periods or transcriptional inhibition shows further reduction in nascent rRNA synthesis, which progressively remodels the nucleoli, to reorganize its proteins and compartments, gradually disrupting the RNA protein interactions within the nucleoli which was essential for the integrity of the tripartite structure. Eventually the compartments form the nucleolar caps observed by a shift from positive to negative scoring of the distance metric between UBF1 foci and the nucleolar periphery (Figure 5B). Several GC-resident proteins like NCL and NPM shuttle out to nucleoplasm to aid in the DNA repair process by p53 dependent pathway(Al-Baker *et al*., 2004; Daniely *et al*., 2002b; Yang *et al*., 2016). There is concomitant reduction in rRNA synthesis which in turn leads to increased protein-protein interaction due to higher affinity and missing rRNA which generally scaffolds multivalent interaction among different types of proteins. We thus observed a liquid to solid-like transition in one of the sub-nucleolar condensates as evidenced by reduced GFP-UBF dynamics (Figure 6A). The GFP-UBF foci show reduced recovery upon total loss of rRNA with direct RNAPI inhibition by BMH21 (Figure 6B, C). The 4NQO doses still showed residual rRNA and hence far-less pronounced reductions in GFP-UBF1 dynamics. Concomitantly the pH gradient within the nucleolus thus shifts towards higher pH values due to reduced negative charge from synthesizing rRNA (Figure 6D, E). Interestingly 4NQO shows a more pronounced effect here than BMH21. This may not be unreasonable, as there are determinants of the pH gradient beyond just rRNA. The alteration in pH is due to change in molecular association underlying the phase-separation of nucleoli due to difference in local protein concentration, ionic strength, and hydrogen bonding(Detres *et al*, 2025; Dogra & Kriwacki, 2025). A recent study has shown that one of the chemical signaling components responsible for the deacidification of the nucleolus upon stress or transcriptional decline, is Poly-ADP-ribose-Polymerase1 (PARP1)(Aryan *et al*., 2023). It is seen that in response to stress nucleolar PARP1 pool delocalizes to the nucleoplasm. The enzymatic activity of PARP1 causes poly-ADP-ribosylation (PARylation) of multiple nucleolar proteins which help maintain nucleolar scaffold during the rRNA synthesis. This is perturbed during stress, leading to change in the proton gradient and altering the pH of the nucleoli(Aryan *et al*., 2023). PAR chains are negatively charged and loss of nucleolar pools and nucleoplasmic recruitment of PARP1 with DNA damage, can cause more pronounced deacidification with loss of both nascent rRNA and PAR. DNA damage also causes the translocation of nucleolar proteins rich in DE tracts(Feric *et al*., 2016; Yang *et al*., 2016). Together nascent rRNA reduction and alteration of the biophysical properties of the nucleolus, marks initiation of formation of nucleolar caps, as the rRNA now gradually forms a hemi ring due to reduced available networks (Figure 4A). Finally, the cap formation is complete when persistent damage or RNAPI inhibition leads to significant reduction in nascent transcripts and the nucleolar proteins forming the FC-DFC (UBF1, and FIB in this study). This eventually leads to complete transformation of the viscoelastic sub-organelle, to a low dynamics multi-phase condensate with increased solid-like properties in the FCs (Figure 5A-C). Altogether this study highlights how the ongoing nascent rRNA synthesis regulate the physical and chemical properties of the multiphase nucleolus and its reorganization upon stress (Figure 6) which can eventually leads to repair of damaged DNA lesions within the nucleoli, if any. So instead of active DNA damage signaling for rDNA repair, nucleolar reorganization can arise naturally from the biophysical changes to the nucleolus under stress. Given that nucleolus plays a critical role in health and diseases, further investigation of how individual segments of rRNA, nascent or mature, and other non-coding RNA (modulating rRNA modification processes and ribosome assembly), or nucleolar PARylation and pH gradients, can affect the physicochemical properties of the sub-organelle will be important for fundamental understanding of nucleolar function.

## Materials and methods

### Cell culture and treatments

U2OS cells were grown in McCoy 5A media (Sigma, M4892) supplemented with 10% FBS (Gibco, 16000-044) and 1% PenStrep-glutamine (Gibco, 10378-016). Cells were grown in T-25 flasks (Tarsons, 950040) and incubated in a CO_2_ incubator (Eppendorf Galaxy, 170S) set to 37℃ and 5% CO_2_. For experiments and imaging, the cells were plated in a glass-bottom dish (Genetix, 200350) at least 24 hours before the experiments. For live cell experiments, the cells were imaged directly in a supplemented FluoroBrite medium (Gibco, A1896701). The plates were placed in the live-cell chamber and maintained at 37°C. The cells are equilibrated for 15 minutes prior to imaging. Transfection was performed using the X-tremeGENE™ HP DNA Transfection Reagent (Roche, 6366236001) following the manufacturer’s protocol. For inhibition of RNA Polymerase-I, BMH21 (ab219384) was used at 2uM for 3 hours. For global DNA damage to the cells 4-nitroquinoline-N-oxide (4-NQO) was used at 1ug/ml for 1 hour or 3 hours. For labeling the RNA, the cells were pulse labeled with 5-Ethynyl uridine (5-EU, CLK-N002-10, Jena Biosciences) for 1mM for 30 minutes and then click chemistry was performed to visualize them. The protocol was adapted from previously described protocols(Jao & Salic, 2008; Loschberger *et al*, 2014). Briefly, after fixation and permeabilization the cells were washed with Tris-buffered saline (TBS) buffer. The cells were then incubated with click reaction cocktail (CRC) for 30 minutes. A 500 µl solution of CRC was prepared as follows: 400 µl of 100 mM Tris (pH ∼8, T6066, Sigma), 10 µl of 100 mM CuSO_4_ (C1297, Sigma), 39 µl of 50mM THPTA (tris-hydroxypropyltriazolylmethylamine) (762342, Sigma), 1 µl of 6.2 mM azide dye (Tamra azide, A7130, Lumiprobe), 50 µl of 1 M sodium ascorbate (11140, Sigma). The cells are then washed with PBS-EDTA (ethylenediaminetetraacetic acid, E6758, Sigma 10mM with 1X PBS) for 10 minutes and followed by two washes with 1X PBS, and immunofluorescence assay can be continued as mentioned below.

### Immunofluorescence (IF)

The cells were fixed in 4% paraformaldehyde (PFA, Sigma P6148) in 1X PBS (phosphate buffered saline, Sigma P5493) for 20 minutes at room temperature. The cells were washed twice with 1X PBS, followed by permeabilization with 0.3% Triton X -100 (Sigma, T8787) in 1X PBS for 10 minutes at room temperature. The cells were washed twice with 1X PBS and blocked with 5% Bovine serum albumin (BSA, Himedia TC545) for an hour. The cells were incubated with the primary antibodies in 5% BSA, at 4℃ overnight. After two washes in 1X PBS, the cells were incubated with the secondary antibodies in 5% BSA, for 2 hours, at room temperature, in the dark. The cells were washed thrice in 1X PBS and treated with Hoechst (1ug/ml) in 1X PBS for 10 min.

### Gelation, digestion and expansion

For expansion microscopy, we adapted from previously described protocols(Asano *et al*., 2018; Cho *et al*., 2018; Gao *et al*., 2017; Tillberg, 2016; Wassie *et al*., 2019). Cells were grown on coverslips placed in plastic bottom dishes; to fix and stain as mentioned above, the cells were re-fixed with 0.25% glutaraldehyde (G6257, Sigma) for 10 minutes. The biomolecules were then anchored overnight using 0.1mg/ml Acryloyl-X SE (AcX-SE, A20770, Invitrogen) in PBS overnight at room temperature. The cells were washed twice with PBS for 15 minutes each. Meanwhile the gelling components are thawed in ice namely, monomer solution (constituents in table S3), APS (A3678, Sigma) and TEMED (T7024, Sigma) stock solutions. The gelling solution was then made using monomer solution, water, 10% TEMED solution, 10% APS in 47:1:1:1 ratio, in this order and vortex for a few seconds. Immediately after, PBS was removed, the glass coverslips were removed from the plastic bottom dish using forceps. 200μl of gelling solution was added on a parafilm kept in a humidified chamber, and the coverslip was inverted (with the cells facing downwards) on the gelling solution. The chamber was then kept for polymerization for an hour at 37℃.The gel is trimmed using a razor blade. The gel attached to the coverslip is then placed in a 6 well plate with a digestion buffer (2-3 ml, constituents in table S3) prepared beforehand, with freshly added proteinase K (8U/ml). The gel is left immersed in the digestion buffer overnight (12-16h) at room temperature (28-30℃) in the dark. Once the gel got detached from the coverslip post digestion, it was transferred to a petri dish using a paintbrush or another coverslip. The petri dish is filled with water to expand the gel. The water is exchanged thrice every 20 minutes. Excess water was then removed and the gel was cut into smaller pieces and placed on a poly-D lysine coated coverslip for imaging.

### rRNA FISH

For RNA FISH, we adapted from previously described protocols (Dhuppar & Mazumder, 2018, 2020; Mazumder *et al*, 2013; Pasnuri *et al*., 2023). Briefly, cells were fixed with 4% PFA in nuclease-free (NF) PBS for 20 minutes at room temperature, followed by permeabilization with 0.3% Triton X-100 in NF PBS for 10 min at room temperature. The cells were then washed twice with PBS followed by blocking (if combined with IF) using 1% NF BSA and 2% NGS (normal goat serum) for an hour. The cells were then rinsed once each with 1X PBS and wash buffer. The cells were then incubated with a hybridization mix consisting of hybridization buffer (Stellaris), 10% formamide (AM9344, Ambion), RNA FISH probes and primary antibody (optional, if combined with IF), overnight at 37 in a dark, humidified chamber. The next day the cells were washed with wash buffer twice for 30 minutes each at 37, in a humidified chamber. The cells were then briefly washed with 2X SSC and then incubated in secondary antibodies in 2% NGS in 1X PBS at room temperature, in the dark. The cells were then rinsed in 1X PBS, and stained with 1ug/ml Hoechst dye for 10 minutes. Given the high density of rRNA within nucleoli individual rRNA molecules are not distinguished like mRNA in the previous studies. But distinct spatial localization of 5’ETS and 28S are clear. Intensities of each species are measured at a single nucleolar level for quantification.

### pH measurements using SNARF-4F

SNARF-4F (S23921, Invitrogen) stock were made in DMSO, at 5mM. 7.5µM was used in live, confluent U2OS cells seeded on a culture dish, and incubated for 30 minutes. In case of treatments using 4-NQO and BMH21, SNARF-4F was added in the last 30 minutes of their respective incubation times. SNARF-4F was aspirated and the cells were image in Fluorobrite along with the treatment or vehicular control (DMSO). The control plates were imaged first and then the pH calibration was done using pH calibration kit (P35379, Thermo-fisher) according to the manufacturers’ protocol.

### Microscopy and Image acquisition

The images were acquired in Nikon Eclipse Ti2 E, with the additional module of Yokogawa CSU W1 So-Ra, with 60X (Plan-Apo N 60X, NA-1.42, oil). The FRAP images were acquired in 60X oil objective (Plan-Apo N 60X oil, N.A -1.42, oil, Olympus), optical zoom 4.5X, mounted on Olympus IX83 inverted microscope with an attached module for laser scanning confocal - FV 3000 with 488 solid state lasers used at 30% power over a circular ROI of 1 μm^2^, at 1024x1024 pixels. The STED images were acquired using 60 X oil objective (Plan-Apo N 60X oil, N.A -1.42, oil, Olympus IX83) with an attached Abberior STEDYCON system.

For pH measurements, samples were excited using 561nm laser, and emission was collected at 640 (700/75) and 580nm (595/40) using the mentioned filter wheels. Since the Nikon spinning disc had a single camera, the sample was excited twice with the same laser, and the emissions were collected sequentially. The exposure setting for the excitation were 500 ms (em-640) and 3s (em-580). This exposure settings and order of acquisition was kept constant across treatments, experiments and pH calibration.

### Image and data statistical analysis

The line profiles were made in Fiji across single plane images of the respective channels used during imaging. All analyses except the ratio measurements in the nucleolar pH assay, FRAP analysis and intensity measurement of Ki-67were done on MATLAB using custom-made codes. Binary nuclear masks were made using the Hoechst images, while nucleolar masks were extracted from the 28S rRNA images. Nucleoli that were over or under-segmented were eliminated from the analysis. The 5’ETS intensity levels were calculated using the 28S rRNA masks itself, along with the nucleolar area. The nucleolar protein intensities of UBF1 were calculated after automated segmentation of the masks created from the individual images itself since taking the nuclear mask would lead to underestimation of the mean protein levels due their sub-compartmentalization. Circularity was calculated using in-built MATLAB function. Nucleoli per cell was calculated manually.

The Ki-67 intensity levels were manually calculated in Fiji for a single plane, with freehand marking of the nucleoli.

For calculating the ring-like structure of 5’ETS and the UBF, binary masks were created from UBF images and annulus mask was created around the UBF mask. The intensity of UBF was calculated within the foci and at the annulus. Similarly,5’ETS intensity in the foci and the annulus was also calculated similarly, and its ratio was plotted.

The nucleolar reorganization was quantified using a metric we designed. The cell nucleoli from the most focused plane were manually segmented from the Hoechst image, and their perimeter was marked, and perimeter coordinates were extracted. The nascent rRNA intensity levels were measured within the marked nucleoli. The FC centers marked by UBF1 in a nucleolus were segmented using the UBF images itself, and the centroid was calculated for each. The shortest distance between the individual UBF1 foci and the nucleolar periphery was then calculated. The distance thus calculated from the UBF centroid to the nucleolar periphery was annotated as positive if the foci was within the nucleolar perimeter and negative if the centroid of the foci lay outside the nucleolar periphery. The nascent rRNA levels is then plotted against the average distance of the UBF1 foci from the nucleolar periphery of a given nucleolus.

For FRAP analysis, a constant circular ROI is used to mark the photobleached region and the intensity was calculated for each timepoint before, and after bleaching. The entire nucleus was marked with freehand ROI and its intensity was also calculated similar to the bleached region. A constant background (bg) value is subtracted from the photobleached ROI (ROI-bg = I) and also from the nucleus (Nuc-bg = total). Then the FRAP index was calculated as follows: (I_(t)_/I_(0)_ * total _(t)_/total_(0)_): I_(t)_ is intensity of the bleached ROI at each timepoint, I_(0)_ is the intensity of the ROI before bleaching, total _(t)_ is the intensity of the nucleus at each timepoint and total_(0)_ is the intensity of the nucleus before bleaching).

The 3D renders of the FCs marked by UBF in the expansion microscopy experiments were obtained using IMARIS software.

For the pH measurements, individual channels were taken after constant background subtraction and manual marking of the nucleoli was done using the freehand tool, on Cy5 channel which showed clear nucleolar signal, and the intensities were recorded using Fiji. All graphs and statistical measurements were made on Origin. To compare means, quantification was done using data from at least three independent replicates unless mentioned otherwise. The bar graphs represent the mean and standard error of the mean (SEM) unless mentioned otherwise. For single-cell distributions, a non-parametric test (Kolmogorov-Smirnov test) was employed as it does not make assumptions regarding the normality of the underlying data. Single-cell distributions may not follow a normal distribution, and a non-parametric test is more suited for these cases. p values below 0.05 were considered statistically significant.

### Antibodies, FISH probes, and chemicals used

The dilutions of primary and secondary antibodies used for IF are given in Table S1. For certain antibodies a higher concentration was used for expansion microscopy experiments. GFP-tagged UBF was obtained from Addgene (Addgene plasmid #17656). Probe sequences targeting rRNA were procured from Pixel Biotech GmBH with required fluorophores (Atto-488 for 5’ETS probes or Atto-565 for 28S probes). The probe sequences are presented in Table S2. Multiple probes were used against each target sequence. Each probe carried a single fluorophore. Chemicals used are presented in Table S3.

---

**Table S1.**

| **Antibody** | **Dilution** | **Catalogue number** |
| --- | --- | --- |
| Rabbit UBF1 | 1:1000 / 1:500 | ab24487-Abcam |
| Rabbit UBF1 | 1:1000/ 1:500 | PA5-82245-Invitrogen |
| Rabbit NPM1 | 1:1000/1:500 | ab183340/15440, Abcam |
| Rabbit POLR1A | 1:1000/1:500 | ab222065-Abcam |
| Mouse NCL | 1:1000/1:500 | E5M7K, CST |
| Chicken FIB | 1:500 | PA5-143565- Invitrogen |
| Rabbit Ki-67 | 1:500 | ab16667 - Abcam |
| Rabbit secondary 488 | 1:1000/1:500 | A11034-Thermo |
| Rabbit secondary 568 | 1:1000/1:500 | A11011-Thermo |
| Rabbit secondary 647 | 1:1000/1:500 | A21245- Thermo |
| Rabbit secondary atto 647 | 1:500 | 40839, CST |
| Mouse secondary 488 | 1:1000/1:500 | A11029-Thermo |
| Mouse secondary atto 647 | 1:500 | 50185, CST |
| Chicken secondary 488 | 1:1000/1:500 | A11039-Thermo |
| Chicken secondary 647 | 1:1000 | A21449- Thermo |

**Table S2.**
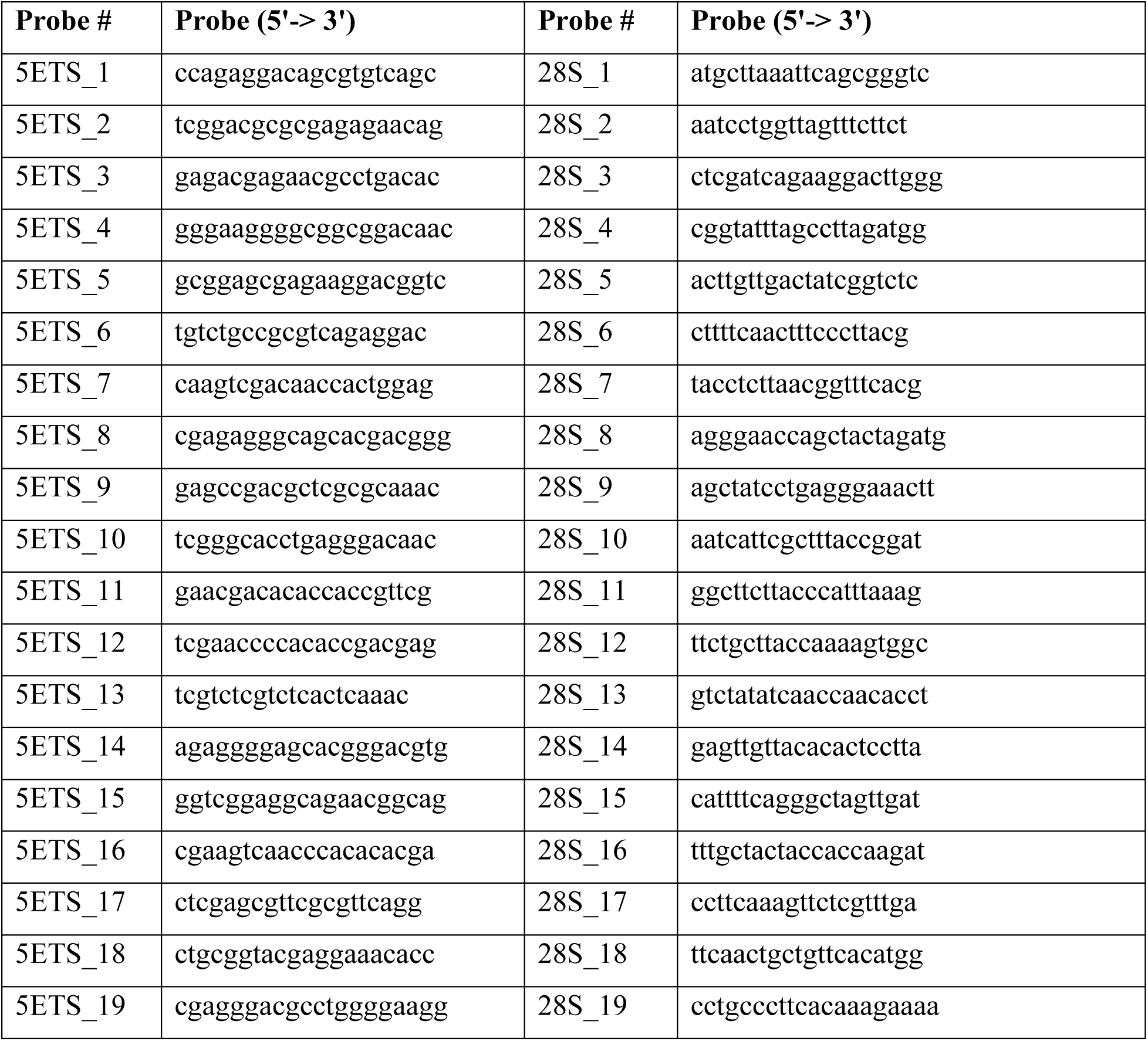
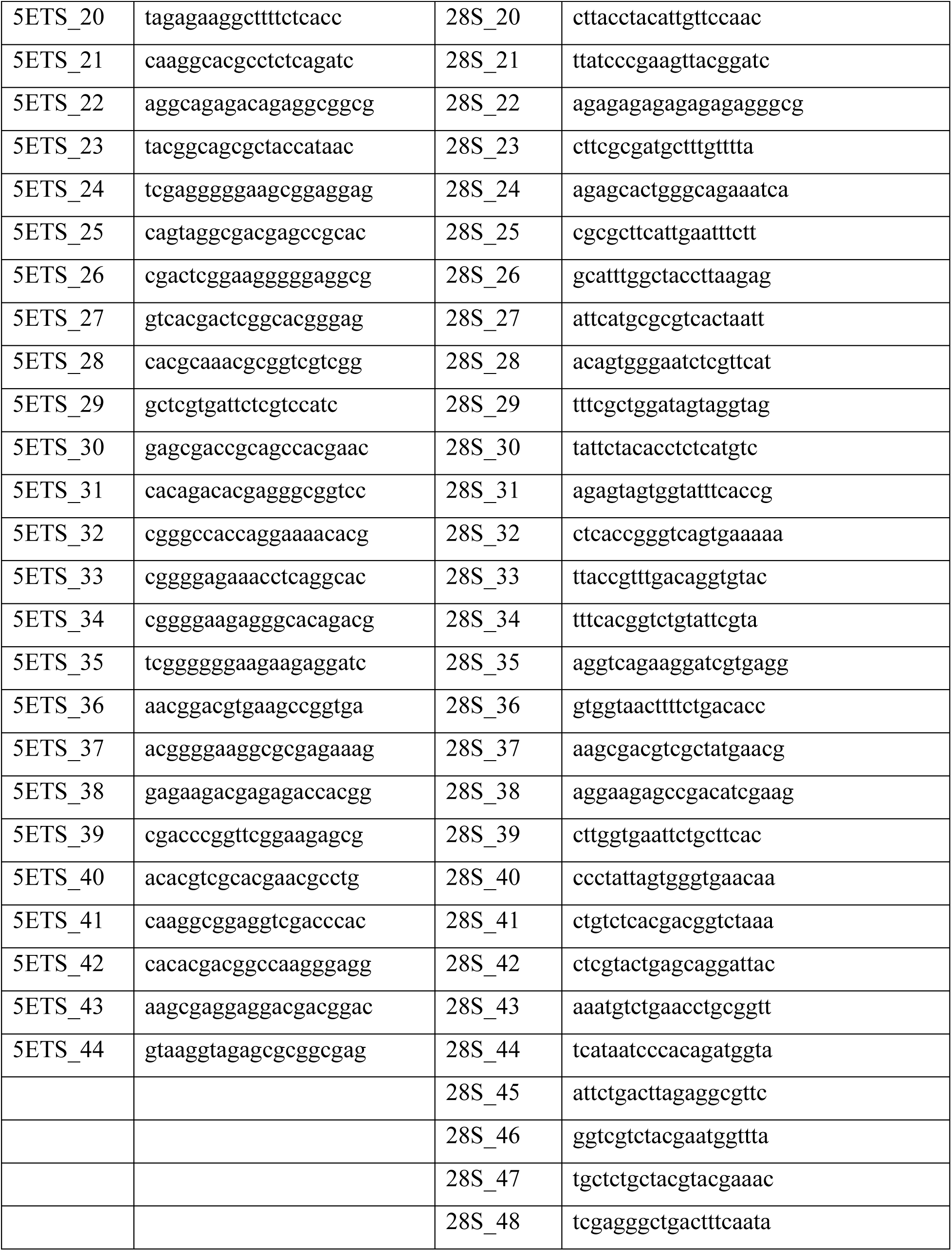
Table with custom-made probe sequences targeting 5’ETS rRNA (44 20 -nucleotide DNA oligos) or 28S mature rRNA (48 20 -nucleotide DNA oligos).

**Table S3.**

| **Monomer Solution** | **Stock Solution** | **Final concentration(g/100ml)** | **Catalogue Number** |
| --- | --- | --- | --- |
| Sodium Acrylate | 38g/100ml | 8.6 | 408220, Sigma |
| Acrylamide | 40g/100ml | 2.5 | A4058, Sigma |
| N,N-Methylenebisacrylamide | 2g/100ml | 0.15 | M1533, Sigma |
| Sodium chloride | 5M | 11.7 | 746398, Sigma |
| PBS | 10X | 1X | P5493 |
| Water |  |  |  |
| **Digestion buffer** | **Stock solution** | **Final concentration(g/100ml)** | **Catalogue number** |
| Triton X-100 | - | 0.50g | T8787, Sigma |
| EDTA | 0.5M, pH-8 | 0.2ml | E6758, Sigma |
| Tris-Cl | 1M, pH-8 | 5ml | T6066, Sigma |
| Sodium chloride | - | 4.67g | 746398, Sigma |
| Water | - | - |  |
| Proteinase-K | 800U/ml | 8U/ml | P2308, Sigma |

## Acknowledgements

This project was supported by intramural funds of TIFR Hyderabad from the Department of Atomic Energy, Government of India (Project Identification No. RTI 4007) GFP-UBF was a gift from Tom Misteli (Addgene plasmid #17656; RRID: Addgene 17656). We thank DSS Imagetech Pvt. Ltd., New Delhi, India, for arranging the demonstration of the Abberior STEDYCON super-resolution microscopy system and for their technical assistance during the imaging process. We thank Purnati Khuntia for kindly sharing access to the IMARIS software (licensed B7y4-du9y-5k5j-76bn) which helped in making the 3D renders used in this study. We thank the Imaging and Flow Facility at TIFR, for their support access to instrumentation.

## Conflict of interest

The authors declare no conflict of interest.

## Notes

### Competing Interest Statement

The authors have declared no competing interest.

## References

Al-Baker EA, Boyle J, Harry R, Kill IR (2004) A p53-independent pathway regulates nucleolar segregation and antigen translocation in response to DNA damage induced by UV irradiation. Exp Cell Res 292: 179–186

Alley KR, Wyatt KM, Fries AC, DeRose VJ (2025) Expansion Microscopy Provides Nanoscale Insight into Nucleolar Reorganization and Nuclear Foci Formation during Nucleolar Stress. ACS Chem Biol 20: 1232–1246

Aryan F, Detres D, Luo CC, Kim SX, Shah AN, Bartusel M, Flynn RA, Calo E (2023) Nucleolus activity-dependent recruitment and biomolecular condensation by pH sensing. Mol Cell 83: 4413–4423 e4410

Asano SM, Gao R, Wassie AT, Tillberg PW, Chen F, Boyden ES (2018) Expansion Microscopy: Protocols for Imaging Proteins and RNA in Cells and Tissues. Curr Protoc Cell Biol 80: e56

Aubert M, O’Donohue MF, Lebaron S, Gleizes PE (2018) Pre-Ribosomal RNA Processing in Human Cells: From Mechanisms to Congenital Diseases. Biomolecules 8

Benjamin R. Sabari^1^, *, Alessandra Dall’Agnese1, Richard A. Young1, (2020) Biomolecular Condensates in the Nucleus. Trends Biochem Sci 45(11): 961–977

Berry J, Weber SC, Vaidya N, Haataja M, Brangwynne CP (2015) RNA transcription modulates phase transition-driven nuclear body assembly. Proceedings of the National Academy of Sciences 112: E5237–E5245

Boamah EK, Kotova E, Garabedian M, Jarnik M, Tulin AV (2012) Poly(ADP-Ribose) polymerase 1 (PARP-1) regulates ribosomal biogenesis in Drosophila nucleoli. PLoS Genet 8: e1002442

Boulon S, Westman BJ, Hutten S, Boisvert FM, Lamond AI (2010) The nucleolus under stress. Mol Cell 40: 216–227

Brangwynne CP, Mitchison TJ, Hyman AA (2011) Active liquid-like behavior of nucleoli determines their size and shape in Xenopus laevis oocytes. Proc Natl Acad Sci U S A 108: 4334–4339

Cahoon CK, Yu Z, Wang Y, Guo F, Unruh JR, Slaughter BD, Hawley RS (2017) Superresolution expansion microscopy reveals the three-dimensional organization of the Drosophila synaptonemal complex. Proc Natl Acad Sci U S A 114: E6857–E6866

Caragine CM, Haley SC, Zidovska A (2019) Nucleolar dynamics and interactions with nucleoplasm in living cells. Elife 8

Carvalho LS, Gonçalves N, Fonseca NA, Moreira JN (2021) Cancer Stem Cells and Nucleolin as Drivers of Carcinogenesis. Pharmaceuticals 14: 60

Chen D, Huang S (2001) Nucleolar components involved in ribosome biogenesis cycle between the nucleolus and nucleoplasm in interphase cells. J Cell Biol 153: 169–176

Cho I, Seo JY, Chang J (2018) Expansion microscopy. J Microsc

Correll CC, Rudloff U, Schmit JD, Ball DA, Karpova TS, Balzer E, Dundr M (2024) Crossing boundaries of light microscopy resolution discerns novel assemblies in the nucleolus. Histochem Cell Biol 162: 161–183

Daniely Y, Dimitrova DD, Borowiec JA (2002a) Stress-dependent nucleolin mobilization mediated by p53-nucleolin complex formation. Mol Cell Biol 22: 6014–6022

Daniely Y, Dimitrova DD, Borowiec JA (2002b) Stress-Dependent Nucleolin Mobilization Mediated by p53-Nucleolin Complex Formation. Molecular and Cellular Biology 22: 6014–6022

Detres D, Camacho-Badillo A, Calo E (2025) A pH-Centric Model of Nucleolar Activity and Regulation. J Mol Biol 437: 169136

Dhuppar S, Mazumder A (2018) Measuring cell cycle-dependent DNA damage responses and p53 regulation on a cell-by-cell basis from image analysis. Cell Cycle 17: 1358–1371

Dhuppar S, Mazumder A (2020) Investigating cell cycle-dependent gene expression in the context of nuclear architecture at single-allele resolution. J Cell Sci 133

Dogra P, Kriwacki RW (2025) Phase separation via protein-protein and protein-RNA networks coordinates ribosome assembly in the nucleolus. Biochim Biophys Acta Gen Subj 1869: 130835

Dundr M, Hoffmann-Rohrer U, Hu Q, Grummt I, Rothblum LI, Phair RD, Misteli T (2002) A kinetic framework for a mammalian RNA polymerase in vivo. Science 298: 1623–1626

Farley-Barnes KI, McCann KL, Ogawa LM, Merkel J, Surovtseva YV, Baserga SJ (2018) Diverse Regulators of Human Ribosome Biogenesis Discovered by Changes in Nucleolar Number. Cell Rep 22: 1923–1934

Farley KI, Surovtseva Y, Merkel J, Baserga SJ (2015) Determinants of mammalian nucleolar architecture. Chromosoma 124: 323–331

Femino AM, Fay FS, Fogarty K, Singer RH (1998) Visualization of single RNA transcripts in situ. Science 280: 585–590

Feric M, Vaidya N, Harmon TS, Mitrea DM, Zhu L, Richardson TM, Kriwacki RW, Pappu RV, Brangwynne CP (2016) Coexisting Liquid Phases Underlie Nucleolar Subcompartments. Cell 165: 1686–1697

Franek M, Kovarikova A, Bartova E, Kozubek S (2016) Nucleolar Reorganization Upon Site-Specific Double-Strand Break Induction. J Histochem Cytochem 64: 669–686

Gao R, Asano SM, Boyden ES (2017) Q&A: Expansion microscopy. BMC Biol 15: 50

Halpern AR, Alas GCM, Chozinski TJ, Paredez AR, Vaughan JC (2017) Hybrid Structured Illumination Expansion Microscopy Reveals Microbial Cytoskeleton Organization. ACS Nano 11: 12677–12686

Hernandez-Verdun D (2006a) The nucleolus: a model for the organization of nuc. Histochem Cell Biol 126:135–148

Hernandez-Verdun D (2006b) The nucleolus: a model for the organization of nuclear functions. Histochem Cell Biol 126: 135–148

Jao CY, Salic A (2008) Exploring RNA transcription and turnover in vivo by using click chemistry. Proc Natl Acad Sci U S A 105: 15779–15784

Katelyn R. Alley KMW, Adam C. Fries, and Victoria J. DeRose (2025) Expansion Microscopy Provides Nanoscale Insight into Nucleolar Reorganization and Nuclear Foci Formation during Nucleolar Stress. ACS chemical biology 20: 1232−1246

King MR, Ruff KM, Lin AZ, Pant A, Farag M, Lalmansingh JM, Wu T, Fossat MJ, Ouyang W, Lew MD et al. (2024) Macromolecular condensation organizes nucleolar sub-phases to set up a pH gradient. Cell 187: 1889–1906 e1824

Kobayashi J, Fujimoto H, Sato J, Hayashi I, Burma S, Matsuura S, Chen DJ, Komatsu K (2012) Nucleolin Participates in DNA Double-Strand Break-Induced Damage Response through MDC1-Dependent Pathway. PLoS ONE 7: e49245

Korsholm LM, Gal Z, Nieto B, Quevedo O, Boukoura S, Lund CC, Larsen DH (2020) Recent advances in the nucleolar responses to DNA double-strand breaks. Nucleic Acids Res 48: 9449–9461

Lafita-Navarro MC, Conacci-Sorrell M (2023) Nucleolar stress: From development to cancer. Semin Cell Dev Biol 136: 64–74

Lafontaine DLJ, Riback JA, Bascetin R, Brangwynne CP (2021) The nucleolus as a multiphase liquid condensate. Nat Rev Mol Cell Biol 22: 165–182

Lawrimore J, Kolbin D, Stanton J, Khan M, de Larminat SC, Lawrimore C, Yeh E, Bloom K (2021) The rDNA is biomolecular condensate formed by polymer-polymer phase separation and is sequestered in the nucleolus by transcription and R-loops. Nucleic Acids Res 49: 4586–4598

Lian Zhua TMR, Ludivine Wacheulc, Ming-Tzo Weia, Marina Ferica, Gena Whitneya, Denis L. J. Lafontainec, and Clifford P. Brangwynnea,d,1 (2019) Controlling the material properties and rRNA processing function of the nucleolus. PNAS

Ljungman M (2000) Dial 9-1-1 for p53: mechanisms of p53 activation by cellular stress. Neoplasia 2: 208–225

Loschberger A, Niehorster T, Sauer M (2014) Click chemistry for the conservation of cellular structures and fluorescent proteins: ClickOx. Biotechnol J 9: 693–697

Louvet E, Junera HR, Le Panse S, Hernandez-Verdun D (2005) Dynamics and compartmentation of the nucleolar processing machinery. Exp Cell Res 304: 457–470

Maiser A, Dillinger S, Langst G, Schermelleh L, Leonhardt H, Nemeth A (2020) Super-resolution in situ analysis of active ribosomal DNA chromatin organization in the nucleolus. Sci Rep 10: 7462

Marcotte N, Brouwer AM (2005) Carboxy SNARF-4F as a fluorescent pH probe for ensemble and fluorescence correlation spectroscopies. J Phys Chem B 109: 11819–11828

Martin RM, Ter-Avetisyan G, Herce HD, Ludwig AK, Lattig-Tunnemann G, Cardoso MC (2015) Principles of protein targeting to the nucleolus. Nucleus 6: 314–325

Mazumder A, Tummler K, Bathe M, Samson LD (2013) Single-cell analysis of ribonucleotide reductase transcriptional and translational response to DNA damage. Mol Cell Biol 33: 635–642

Mischo HE, Hemmerich P, Grosse F, Zhang S (2005) Actinomycin D induces histone gamma-H2AX foci and complex formation of gamma-H2AX with Ku70 and nuclear DNA helicase II. J Biol Chem 280: 9586–9594

Mitrea DM, Cika JA, Guy CS, Ban D, Banerjee PR, Stanley CB, Nourse A, Deniz AA, Kriwacki RW (2016) Nucleophosmin integrates within the nucleolus via multi-modal interactions with proteins displaying R-rich linear motifs and rRNA. Elife 5

Pasnuri N, Jaiswal M, Ray K, Mazumder A (2023) Buffered EGFR signaling regulated by spitz-to-argos expression ratio is a critical factor for patterning the Drosophila eye. PLoS Genet 19: e1010622

Pederson T (1998) The plurifunctional nucleolus. Nucleic Acids Res 26: 3871–3876

Penzo M, Montanaro L, Trere D, Derenzini M (2019) The Ribosome Biogenesis-Cancer Connection. Cells 8

Pownall ME, Miao L, Vejnar CE, M’Saad O, Sherrard A, Frederick MA, Benitez MDJ, Boswell CW, Zaret KS, Bewersdorf J et al. (2023) Chromatin expansion microscopy reveals nanoscale organization of transcription and chromatin. Science 381: 92–100

Rai SK, Khanna R, Avni A, Mukhopadhyay S (2023) Heterotypic electrostatic interactions control complex phase separation of tau and prion into multiphasic condensates and co-aggregates. Proc Natl Acad Sci U S A 120: e2216338120

Raj A, van den Bogaard P, Rifkin SA, van Oudenaarden A, Tyagi S (2008) Imaging individual mRNA molecules using multiple singly labeled probes. Nat Methods 5: 877–879

Riback JA, Eeftens JM, Lee DSW, Quinodoz SA, Donlic A, Orlovsky N, Wiesner L, Beckers L, Becker LA, Strom AR et al. (2023) Viscoelasticity and advective flow of RNA underlies nucleolar form and function. Mol Cell 83: 3095–3107 e3099

Roden C, Gladfelter AS (2021) RNA contributions to the form and function of biomolecular condensates. Nat Rev Mol Cell Biol 22: 183–195

Rubbi CP, Milner J (2003) Disruption of the nucleolus mediates stabilization of p53 in response to DNA damage and other stresses. EMBO J 22: 6068–6077

Scott DD, Oeffinger M (2016) Nucleolin and nucleophosmin: nucleolar proteins with multiple functions in DNA repair. Biochem Cell Biol 94: 419–432

Shan L, Xu G, Yao RW, Luan PF, Huang Y, Zhang PH, Pan YH, Zhang L, Gao X, Li Y et al. (2023) Nucleolar URB1 ensures 3’ ETS rRNA removal to prevent exosome surveillance. Nature 615: 526–534

Shav-Tal Y, Blechman J, Darzacq X, Montagna C, Dye BT, Patton JG, Singer RH, Zipori D (2005) Dynamic sorting of nuclear components into distinct nucleolar caps during transcriptional inhibition. Mol Biol Cell 16: 2395–2413

Sheu-Gruttadauria J, Yan X, Stuurman N, Vale RD, Floor SN (2024) Nucleolar dynamics are determined by the ordered assembly of the ribosome. *bioRxiv*

Sofia A. Quinodoz LJ, Aya A. Abu-Alfa, Troy J. Comi, Hongbo Zhao, Qiwei Yu5 LWW, Jordy F. Botello, Anita Donlic, Elizabeth Soehalim, Prashant Bhat, CZ, Ludivine Wacheul, Andrej Košmrlj , Denis L. J. Lafontaine, Sebastian Klinge & Clifford P. Brangwynne (2025) Mapping and engineering RNA-driven architecture of the multiphase nucleolus. nature article

Stenström L, Mahdessian D, Gnann C, Cesnik AJ, Ouyang W, Leonetti MD, Uhlén M, Cuylen-Haering S, Thul PJ, Lundberg E (2020) Mapping the nucleolar proteome reveals a spatiotemporal organization related to intrinsic protein disorder. Molecular Systems Biology 16

Szaflarski W, Lesniczak-Staszak M, Sowinski M, Ojha S, Aulas A, Dave D, Malla S, Anderson P, Ivanov P, Lyons SM (2022) Early rRNA processing is a stress-dependent regulatory event whose inhibition maintains nucleolar integrity. Nucleic Acids Res 50: 1033–1051

Thomas Haaf aDCW ((1996)) Inhibition of RNA Polymerase II Transcription Causes Chromatin Decondensation, Loss of Nucleolar Structure, and Dispersion of Chromosomal Domains. EXPERIMENTAL CELL RESEARCH

Tillberg PW (2016) Protein-retention expansion microscopy of cells and tissues labeled using standard fluorescent protei. Nat Biotechnol 34(9): 987–99

van Sluis M, McStay B (2017) Nucleolar reorganization in response to rDNA damage. Curr Opin Cell Biol 46: 81–86

Vanden Broeck A, Klinge S (2024) Eukaryotic Ribosome Assembly. Annu Rev Biochem 93: 189–210

Wadsworth GM, Srinivasan S, Lai LB, Datta M, Gopalan V, Banerjee PR (2024) RNA-driven phase transitions in biomolecular condensates. Mol Cell 84: 3692–3705

Wassie AT, Zhao Y, Boyden ES (2019) Expansion microscopy: principles and uses in biological research. Nat Methods 16: 33–41

Wei J, Zhang Y, Ye B, Fan X, Hu Y, Xiang S, Ma W (2024) Multivalent 28S rRNA Is the Organizer of the Nucleolus’s Multi-layered Architecture. bioRxiv: 2024.2003.2013.584914

Yamamoto T, Yamazaki T, Ninomiya K, Hirose T (2023) Nascent ribosomal RNA act as surfactant that suppresses growth of fibrillar centers in nucleolus. Communications Biology 6

Yang K, Wang M, Zhao Y, Sun X, Yang Y, Li X, Zhou A, Chu H, Zhou H, Xu J et al. (2016) A redox mechanism underlying nucleolar stress sensing by nucleophosmin. Nat Commun 7: 13599

Yang K, Yang J, Yi J (2018) Nucleolar Stress: hallmarks, sensing mechanism and diseases. Cell Stress 2: 125–140

Yao RW, Xu G, Wang Y, Shan L, Luan PF, Wang Y, Wu M, Yang LZ, Xing YH, Yang L et al. (2019) Nascent Pre-rRNA Sorting via Phase Separation Drives the Assembly of Dense Fibrillar Components in the Human Nucleolus. Mol Cell 76: 767–783 e711

