## Supplementary file for "Nucleolar reorganization on stress depends on physicochemical changes due to nascent rRNA synthesis"

**Authors and affiliations: Sinjini Ghosh[1], and Aprotim Mazumder[1,*]**

1 Tata Institute of Fundamental Research Hyderabad, 36/P, Gopanpally, Serilingampally Mandal, Hyderabad-500046, Telangana, India

Supplementary Figure 1

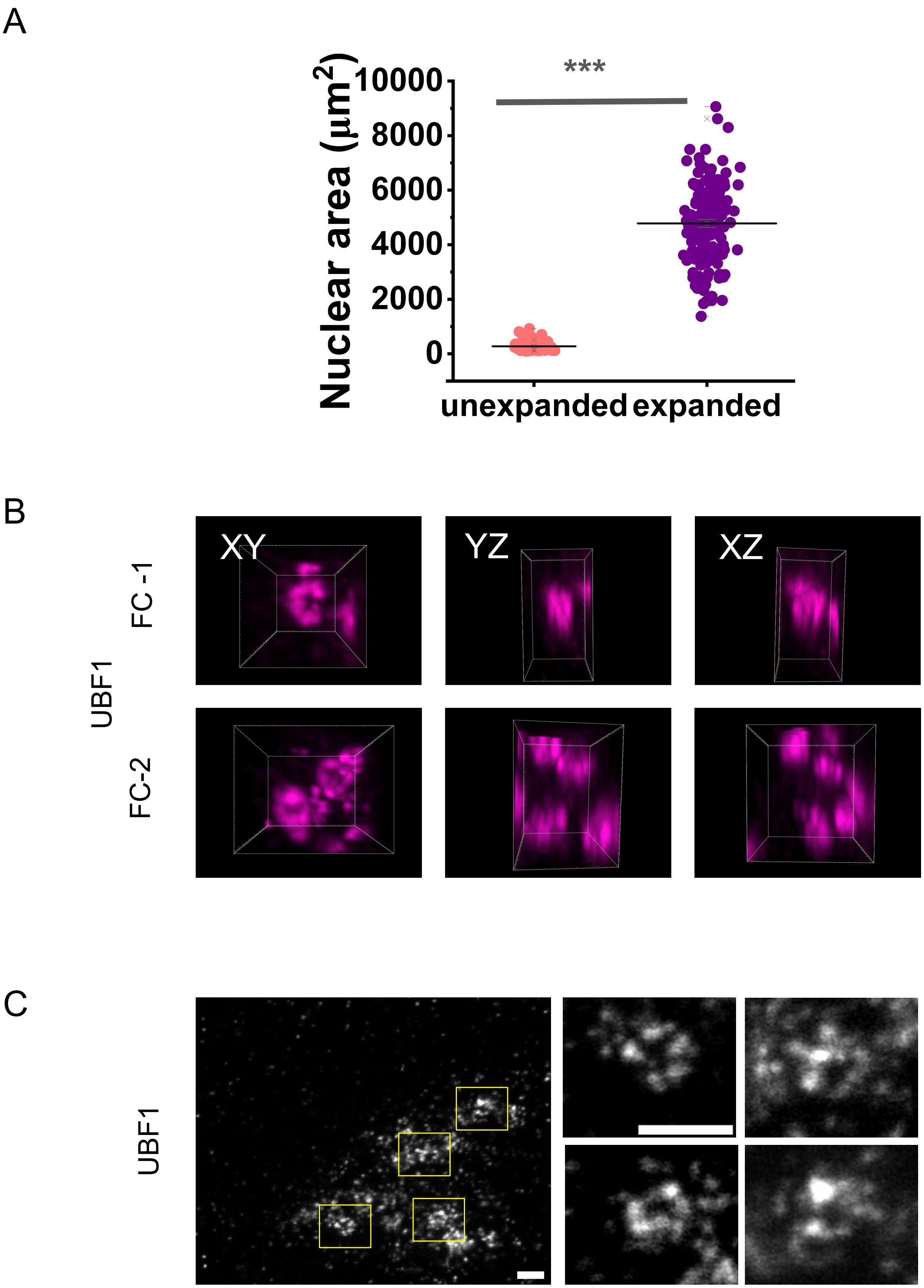

Supp. Figure 1: (A) Graph for nuclear area in unexpanded and expanded nuclei the pink plot.Data represents mean± S.E.M, n=3 for ≥ 60 cells, over three independent experiments. (B) 3D renders of expanded U2OS cell nucleoli showing three different orthogonal slices (left to right ) of XY, YZ, XZ of a 2 FC modules marked by UBF1(magenta). Scale bar is 5 microns. (C) STED images of cells marking UBF1 for a single nucleoli, with 4 FC centres marked in yellow boxes, and the insets shown on its right. Scale bar is 0.5 microns. Statistical significance calculated using K.S. test ( *** represents $p<0.0001$)

Supplementary Figure 2

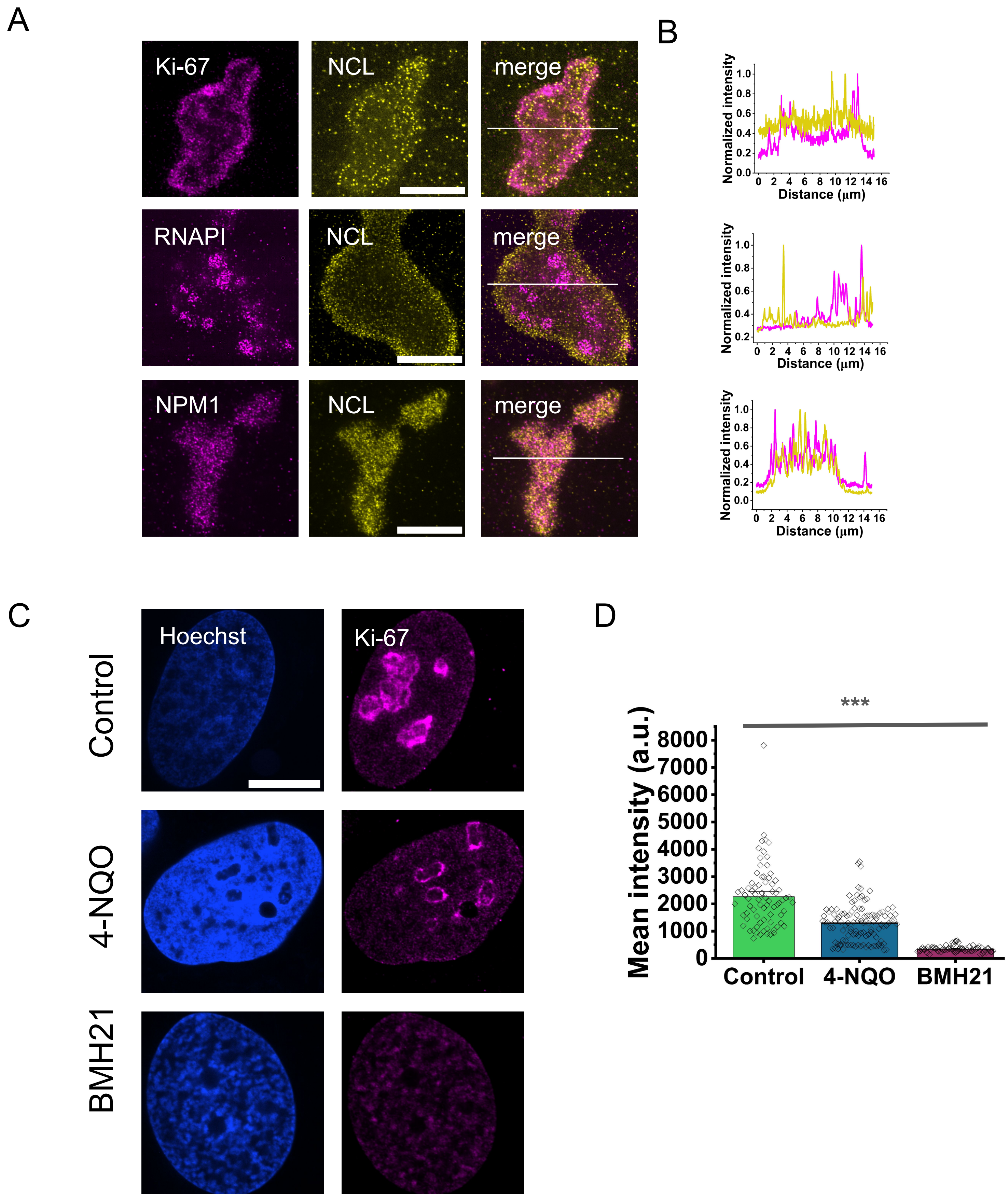

Supp. Figure 2: (A) U2OS cell nucleoli after expansion,marked with Ki-67, RNAP-1, NPM1 (top to bottom respectively, in magenta) co-stained with NCL(yellow) in each case, and merge of the two compartments on the right-most panel. Scale bar is 10 microns. (B) Line profile for the merged images from (A). (C) U2OS cell nuclei treated with 1ug/ml 4NQO for 1hr and 2uM of BMH21 for 3hrs, compared to control nuclei, stained with Ki-67 and its mean nucleolar intensity is plotted in (D), Data represents mean± S.E.M. with n>70, over two independent experiments. Statistical significance calculated using K.S. test (*** represents p<0.0001)

Supplementary Figure 3

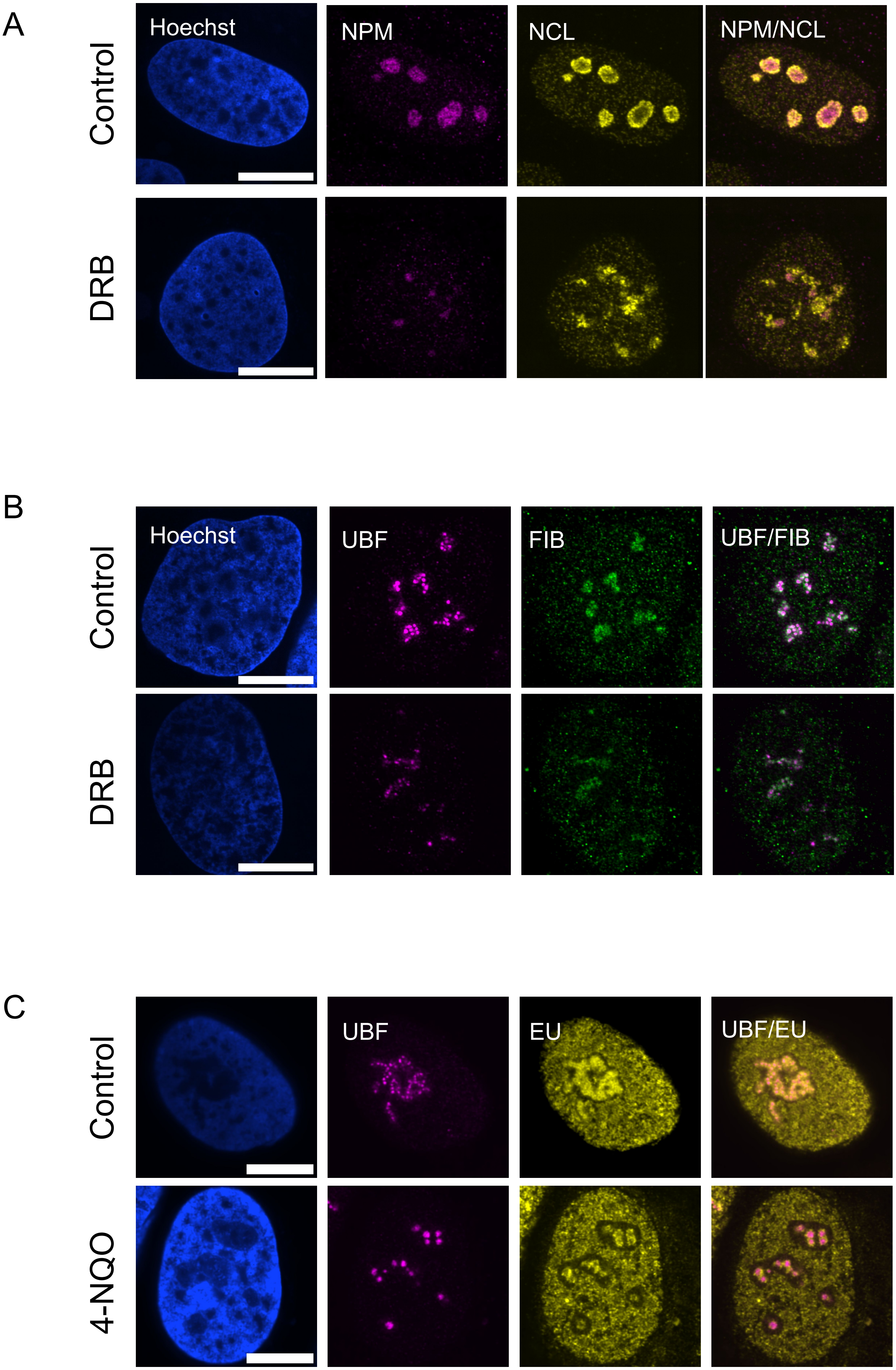

Supp. Figure 3: (A) U2OS cells treated with DMSO (control) and DRB at 60uM, for 1 hour stained with Hoechst (1µg/ml) and nucleoli marked with NPM (magenta), NCL (yellow), and their merge. (B) U2OS cells treated with DMSO (control) and DRB at 60uM, for 1 hour, stained with Hoechst (1µg/ml) and nucleoli marked with UBF (magenta), FIB (green), their merge. (C) shows cells treated with DMSO (control) and 4-NQO at 1ug/ml for 1 hour with 5-EU pulsed for the last 30 minutes to detect nascent transcription. It is then stained with Hoechst (1µg/ml) , with nucleoli marked with UBF1 (magenta) and 5-EU labelled with TAMRA azide (yellow).
